# Across-cohort QC analyses of genome-wide association study summary statistics from complex traits

**DOI:** 10.1101/033787

**Authors:** Guo-Bo Chen, Sang Hong Lee, Matthew R Robinson, Maciej Trzaskowski, Zhi-Xiang Zhu, Thomas W Winkler, Felix R Day, Damien C Croteau-Chonka, Andrew R Wood, Adam E Locke, Zoltán Kutalik, Ruth J F Loos, Timothy M Frayling, Joel N Hirschhorn, Jian Yang, Naomi R Wray, The Genetic Investigation of Anthropometric Traits (GIANT) Consortium, Peter M Visscher

## Abstract

Genome-wide association studies (GWASs) have been successful in discovering replicable SNP-trait associations for many quantitative traits and common diseases in humans. Typically the effect sizes of SNP alleles are very small and this has led to large genome-wide association meta-analyses (GWAMA) to maximize statistical power. A trend towards ever-larger GWAMA is likely to continue, yet dealing with summary statistics from hundreds of cohorts increases logistical and quality control problems, including unknown sample overlap, and these can lead to both false positive and false negative findings. In this study we propose a new set of metrics and visualization tools for GWAMA, using summary statistics from cohort-level GWASs. We proposed a pair of methods in examining the concordance between demographic information and summary statistics. In method I, we use the population genetics *F_st_* statistic to verify the genetic origin of each cohort and their geographic location, and demonstrate using GWAMA data from the GIANT Consortium that geographic locations of cohorts can be recovered and outlier cohorts can be detected. In method II, we conduct principal component analysis based on reported allele frequencies, and is able to recover the ancestral information for each cohort. In addition, we propose a new statistic that uses the reported allelic effect sizes and their standard errors to identify significant sample overlap or heterogeneity between pairs of cohorts. Finally, to quantify unknown sample overlap across all pairs of cohorts we propose a method that uses randomly generated genetic predictors that does not require the sharing of individual-level genotype data and does not breach individual privacy.

## Introduction

Genome-wide association studies (GWASs) have been successful in discovering SNP-trait associations for complex traits^1^. To elucidate genetic architecture, which requires maximized statistical power for discovery of risk alleles of small effect, large genome-wide association meta-analyses (GWAMA) are tending towards ever-larger scale that may contain data from hundreds of cohorts. At the individual cohort level, GWAS analysis is often based on various genotyping chips and conducted with different protocols, such as different software tools and reference populations for imputation, inclusion of study specific covariates and association analyses using different methods and software. Although solid quality control analysis pipelines of GWAMA exist^2^, these analyses focus on quality control (QC) for each cohort independently. With ever-increasing sizes of GWAMA there is a need for additional QC that goes beyond the cohort-by-cohort genotype-level analysis performed to date.

In this study, we propose a new set of QC metrics for GWAMA. In contrast to previous QC metrics, our approach explores the genetic and QC context of the all cohorts in GWAMA together rather than by treating them one at a time. These metrics include

i. a genome-wide comparison of allele frequency differences across cohorts or against a common reference population
ii. principal component analysis for reported allele frequencies
iii. a pairwise cohort statistic that uses allele frequency or effect size concordance to detect the proportion of sample overlap or heterogeneity
iv. an easy to implement analysis to pinpoint each between-cohort overlapping sample that does not require the sharing of individual-level genotype data.

All these applications assume that there is a central analysis hub where summary statistic data from GWAS are uploaded for each cohort. In addition, these metrics reveal information of interest other than merely QC.

## Materials and Methods

### Overview of materials

**Cohort-level summary statistics**. The GWAS height GWAS summary statistics were provided by the GIANT Consortium and were from 82 cohorts (174 separate files due to different ways a cohort was split into different sexes, different disease statuses) representing a total of 253,288 individuals, and nearly 2.5 million autosome SNPs imputed to the HapMap2 reference^3^. The Metabochip summary statistics were for body mass index (BMI) from 43 cohorts (120 files due to different ways a cohort was split into different sexes, different disease statuses) representing a total of 103,047 samples from multiple ethnicities with about 200,000 SNPs genotyped on customised chips^4,5^. For convenience, we consider each file a cohort. All the summary statistics have already been cleaned using established protocols for GWAS meta-analysis^2^.

**1000 Genomes Project samples**. 1000 Genomes Project (1KG) reference samples^6^ were used as the reference samples for calculating *F_st_*. When assessing the global-level *F_st_* measures, Yoruba represent African samples (YRI, 108 individuals), Han Chinese in Beijing represent East Asian samples (CHB, 103 individuals), and Utah Residents with Northern and Western European Ancestry represent European samples (CEU, 99 individuals) were employed as the reference panels. For calculating within-Europe *F_st_*, CEU, Finnish (FIN, 99 individuals), and Tuscani (TSI, 107 individuals) were employed to represent northwest, northeast and southern Europeans, respectively. For analyses using a whole European panel, CEU, FIN, TSI, GBR (British, 91 individuals), and IBS (Iberian, 107 individuals) were pooled together as an “averaged” European reference.

**WTCCC GWAS data**. WTCCC GWAS data has 2,934 shared controls for 7 diseases with a total of 14,000 cases^7^. Individual GWAS was conducted for each disease using PLINK^8^, and their summary statistics used to estimate *λ_meta_* (see text below). WTCCC GWAS data were also used for demonstrating pseudo profile score regression (see text below).

**Simulated cohort-level summary statistics**. *M* independent loci were generated for cohort-level summary statistics. Each locus had allele frequency *p_i_*, which was sampled from a uniform distribution ranging from 0.1 to 0.5, and had genetic effect *b_i_*, sampled from a standard normal distribution *N*(0,1). After rescaling, 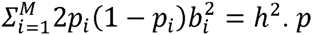 and *b* were treated as true parameters. For a particular cohort with *n* samples, its 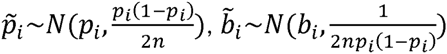, and the sampling variance for 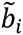 is 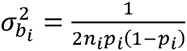. All cohorts were assumed to share common genetic architecture, and differences were only due to genetic drift, allele frequencies and sampling variance of genetic effects.

### Overview of the methods

***F_st_*-based genetic distance between cohorts**. For a cohort, its *F_st_* with reference cohorts, such as CEU, YRI, and CHB, is calculated. Given those three *F_st_* values, the coordinate of this cohort can be uniquely projected into the reference equilateral that has CEU, YRI, and CHB at its corners.

**Principal component analysis for cohort-level allele frequencies**. A genetic relationship matrix for cohorts can be constructed based on received allele frequencies. Principal component analysis (PCA) can be implemented on the genetic relationship matrix. The projection of the cohorts into PCA space can reveal the genetic background and relative geographical distance between cohorts.

λ*_meta_* **for detecting overlapping samples**. In concept, λ*_meta_* resembles *λ_gc_*, which indicates population stratification for a GWAS^9^, but *λ_meta_* measures the proportion of overlapping samples between a pair of cohorts. Based on reported genetic effects and their sampling variance, *λ_meta_* can be constructed for a pair of cohorts and follows a chi-square distribution with 1 degree of freedom. *λ_meta_* will be close to 1 when there is no overlapping samples, smaller than 1 when there are overlapping samples, and greater than 1 when there are heterogeneity between a pair of cohorts. For GWAMA over a single trait across method, we assume heterogeneity is zero.

**Pseudo profile score regression for pinpointing overlapping samples/relatives**. Pseudo profile score regression (PPSR) provides a framework for pinpointing the overlapping samples/relatives between cohort without sharing genotypes. Each GWAS analyst generates pseudo profile scores (PPS) for each sample on a set of loci, which are chosen by a GWAMA central analyst. If the similarity metric of PPS for a pair of cohorts reaches a similarity threshold, say 1 overlapping samples and 0.5 for first-degree relatives, then overlapping samples/relatives are found. PPSR can have a controlled type-I and type-II error rates in pinpointing overlapping samples, and also can reduce the comprise of privacy. PPSR is an enhanced version of Gencrypt^10^, a previous method in pinpointing overlapping samples.

The technical details of these four methods can be found in the Supplementary notes.

## Results

### Population genetic quality control analysis using *F_st_*

Allele frequency differentiation among populations reflects population characteristics such as demographic past and geographic locations^11,12^. In GWAMA only summary statistics such as allele frequencies are available to the central analysis hub, and so it is not possible to run principal component analysis for each cohort that requires individual-level data. Therefore it is difficult to quantify genetic distance between cohorts or to a reference in order to identify population outliers. Outlier cohorts can be due to real differences in ethnicity or mistakes in the primary analysis prior to uploading data to the GWAMA analysis hub. Gross differentiation in allele frequencies at specific SNPs between GWAMA cohorts and a reference (such as 1000 Genomes Project, denoted as 1KG)^6^ are part of standard QC protocols^2^ but checking for more differentiation than expected across the entire genome is not usually part of the QC pipeline. We propose that a genetic distance inferred from *F_st_*, which reflects genetic distance between pairwise populations, is a useful additional QC statistic to detect cohorts that are population outliers. Using the relationship between *F_st_* and principal components^13–15^, our *F_st_* Cartographer algorithm can be used to estimate the relative genetic distance between cohorts (**Supplementary notes, and Supplementary Fig. 1**).

We applied the *F_st_* metric to the GIANT Consortium body mass index (BMI) Metabochip cohorts (55 male-only cohorts, 55 female-only cohort, and 10 mixed-sex cohorts; for convenience, we called each file a cohort), which were recruited from multiple ethnicities^4^, such as Europeans, African Americans in The Atherosclerosis Risk in Communities Study (ARIC) and cohorts from Jamaica (SPT), Pakistan (PROMISE), Philippines (CLHNS) and Seychelles (SEY). For each Metabochip cohort, we sampled 30,000 (see Online method for details) independent markers to calculate *F_st_* values with each of three 1KG samples (CEU, CHB, and YRI, respectively). For validation of the method, we also calculated *F_st_* values against the 1KG Japanese (JPT, Japanese in Tokyo, Japan), Indian (GIH, Gujarati Indian in Houston, US), Kenyan (LWK, Luhya in Webuye, Kenya) and European samples (IBS, Iberian populations, Spain; FIN, Finnish, Finland; TSI, Toscani, Italy, and GBR, British in England and Scortland, GBR), to see whether the known genetic origins of those cohorts can be recovered.

According to the origins of the samples, each Metabochip cohort showed a different genetic distance spectrum to the three reference populations (**Fig. 1a**). The JPT and Philippine cohorts had very small genetic distances to CHB, as expected, but large to CEU and YRI; however, the Pakistan cohorts showed much closer genetic distances to CEU than to CHB and YRI, indicating their demographic history. The cohorts sampled from Jamaica, Seychelles, Hawaii, and the African American ARIC cohort had small genetic distances to YRI, but large distances to CHB and CEU. For most European cohorts, as expected, the distances to CEU were very small compared with those to CHB and YRI. Given their relative distances to CEU, CHB, and YRI, using our *F_st_* cartographer algorithm (**Supplementary notes, and Supplementary Fig. 1**), the cohorts were projected into a two-dimensional space, called *F_st_* derived principal components (*F_PC_*) space, constructed by YRI, CHB, and CEU as the reference populations (**Fig. 1b**). The allocation of the cohorts to the *F_PC_* space resembles that of eigenvector 1 against eigenvector 2 in principal component analysis (PCA)^12^, and is similar to those observed in PCA using individual-level GWAS data for populations of various ethnicities such as in 1KG samples^6^. Therefore, our method to place cohorts in geographical regions from GWAS summary statistics works well at a global-population scale.

**Figure 1.**
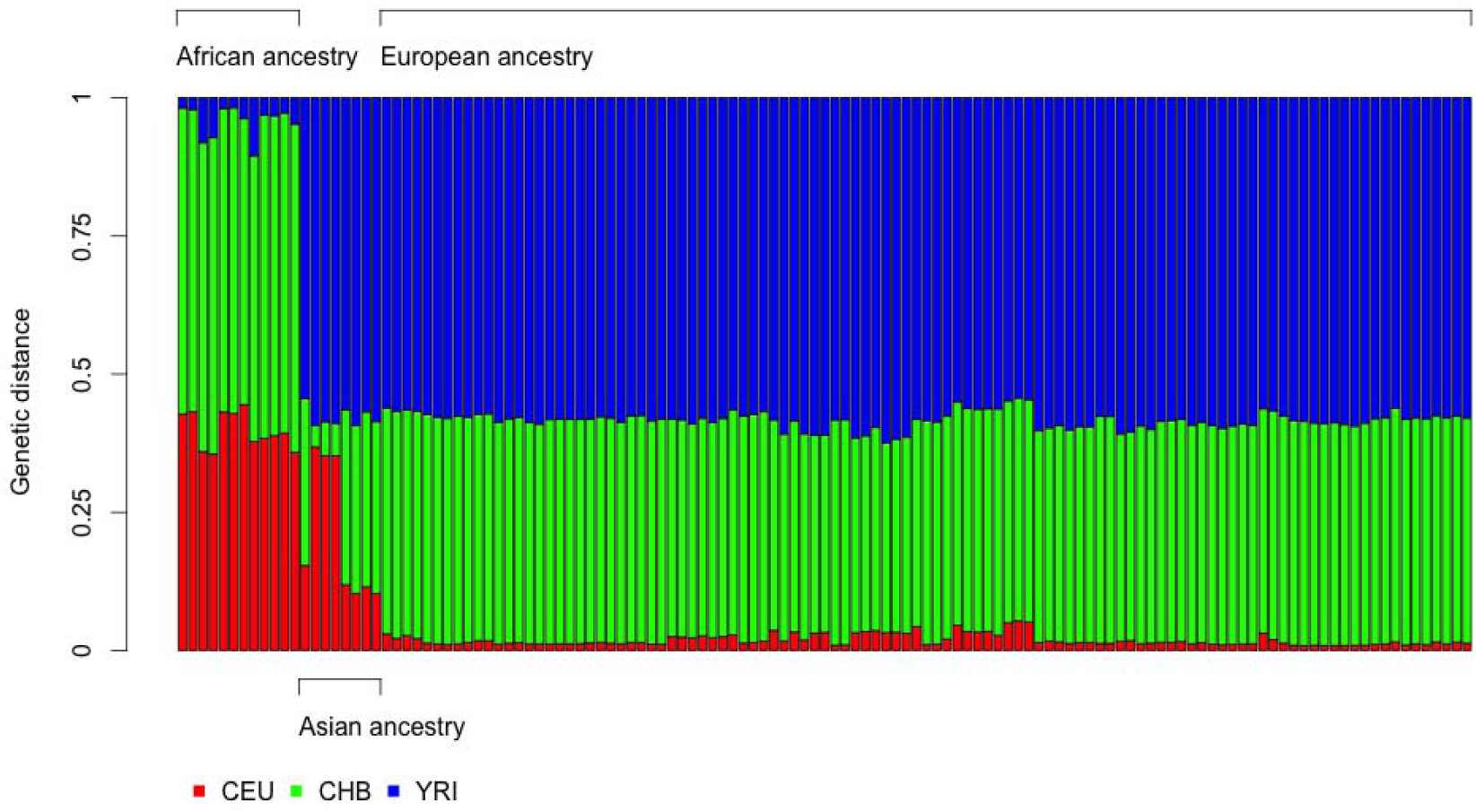
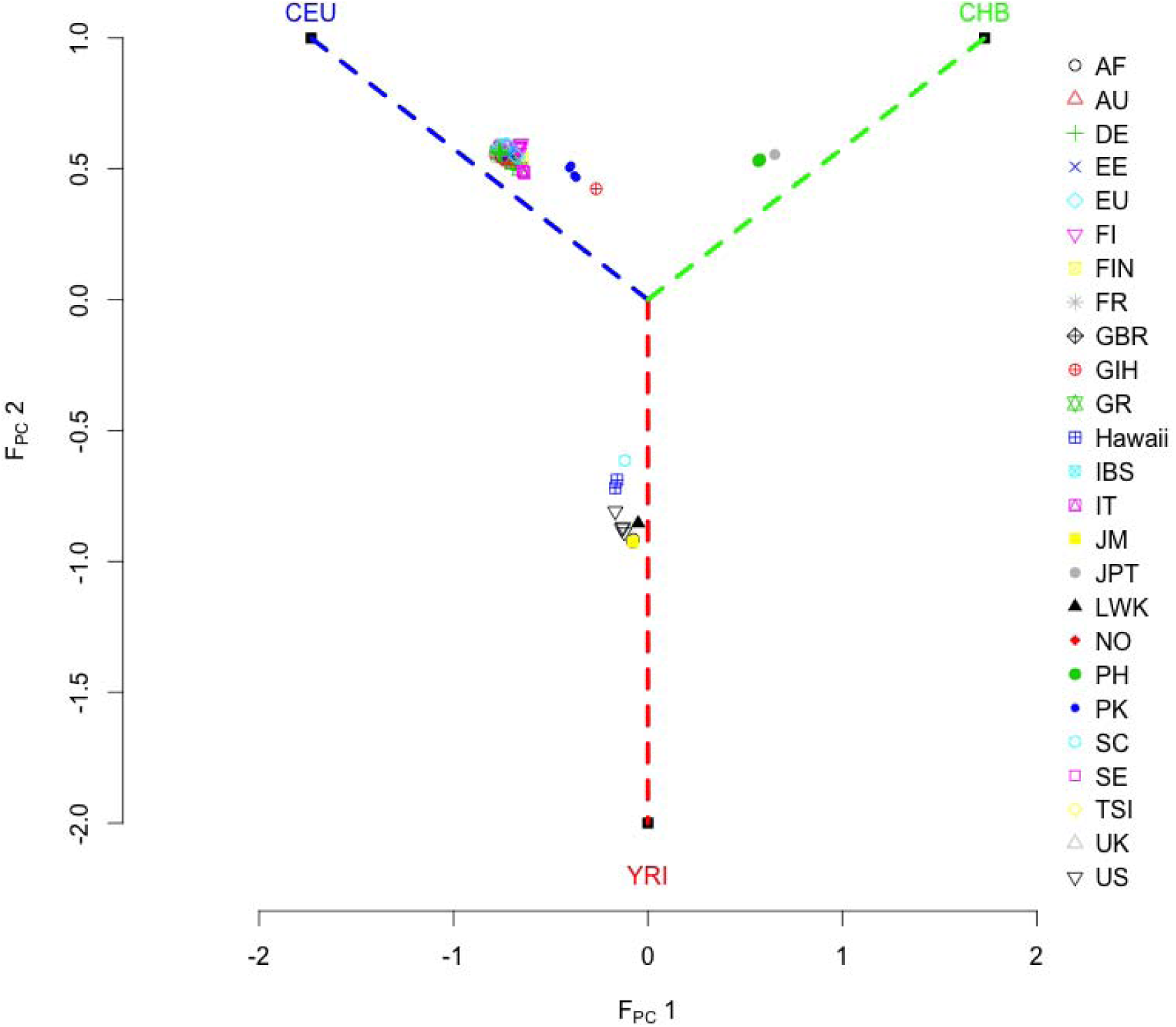
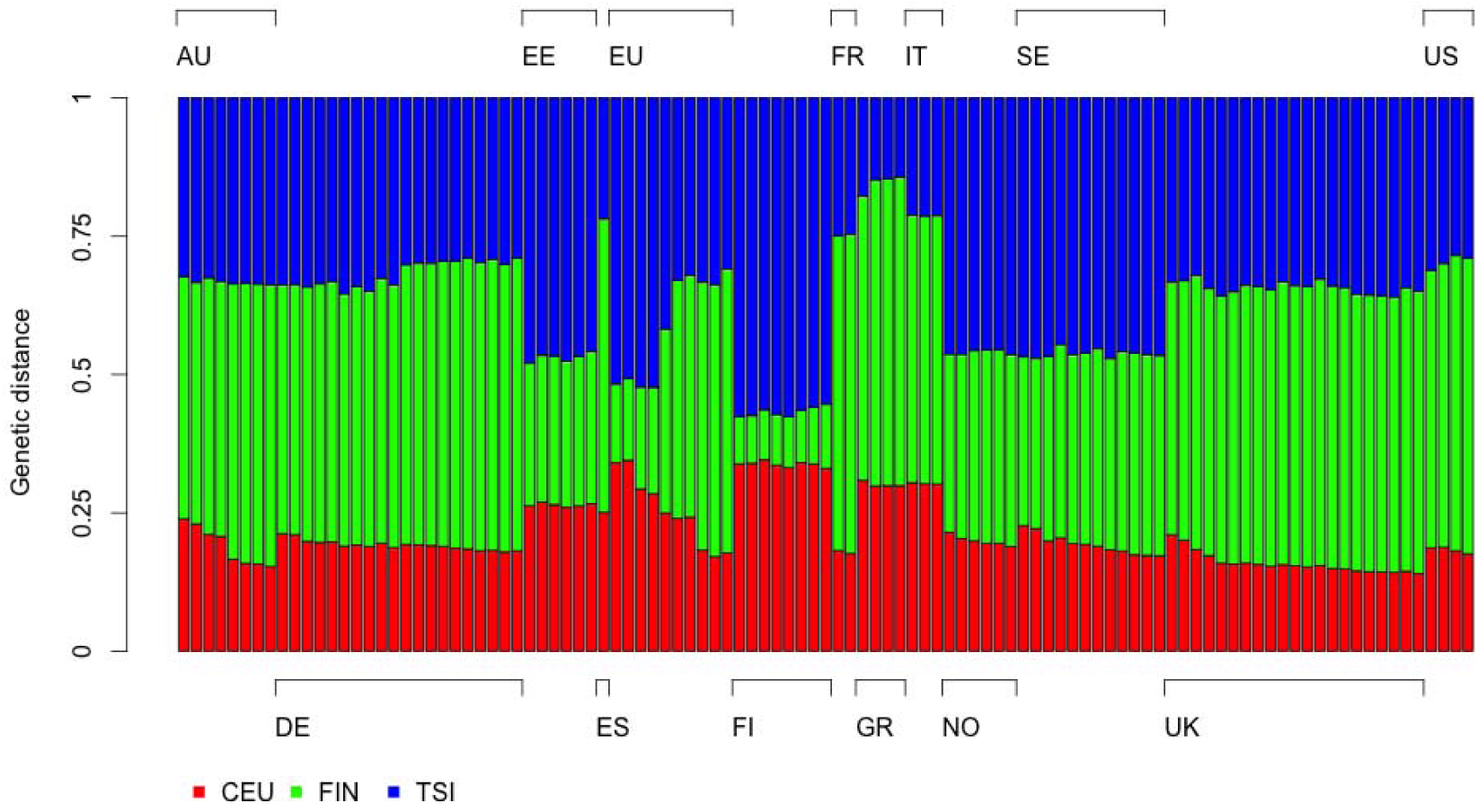
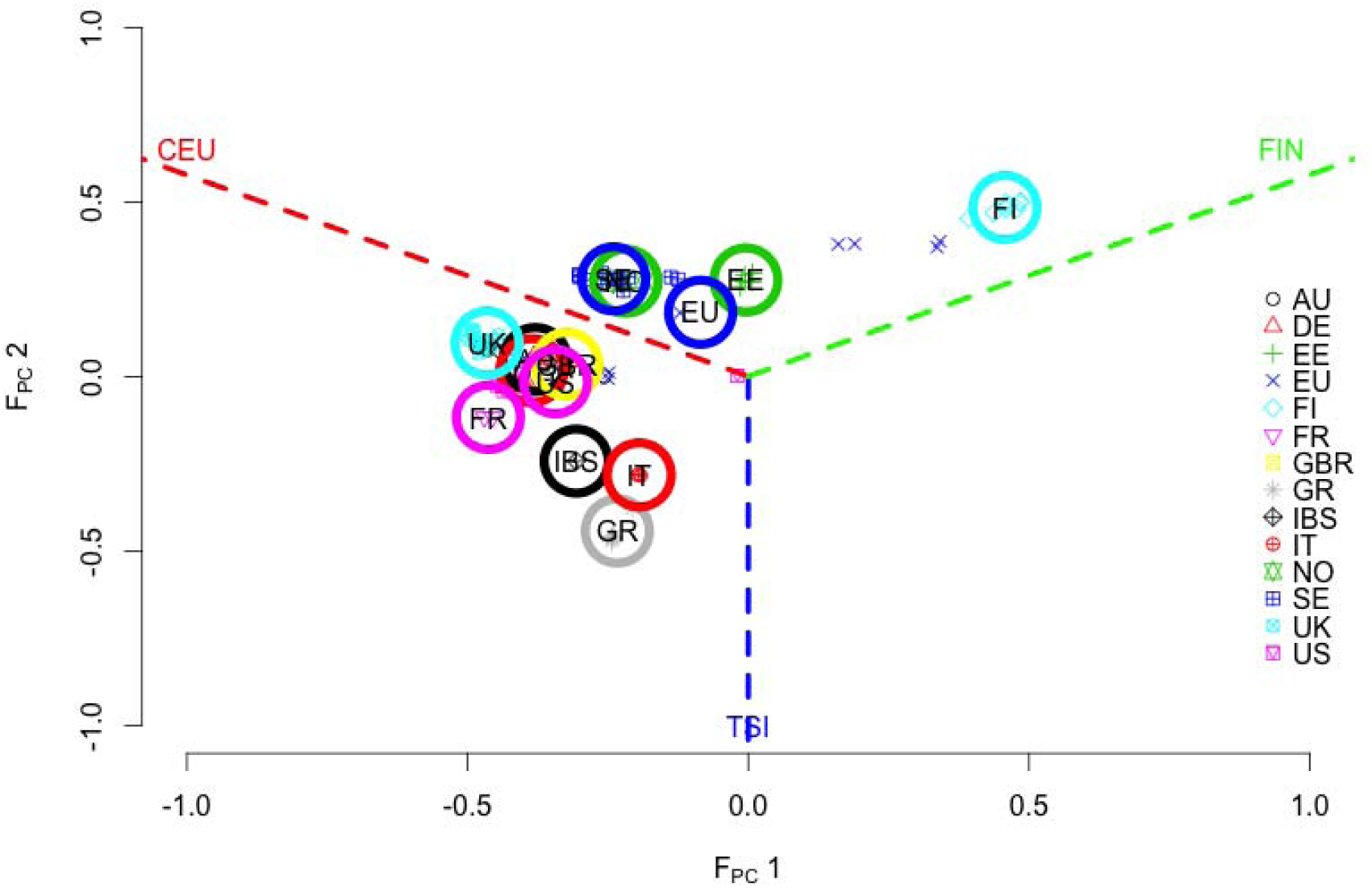
Recovery of cohort-level genetic background and inference of their geographic locations for GIANT BMI Metabochip cohorts using the *F_st_* derived genetic distance measure. **(a)** Genetic distance spectrum for all Metabochip cohorts to CEU, CHB and YRI. See Supplementary notes for more details. The origins of the cohorts are denoted on the horizontal axis. **(b)** Projection for Metabochip cohort into *F_PC_* space defined by YRI, CHB, and CEU reference populations. The x- and y-axis represent relative distances derived from the genetic distance spectrum. Three dashed lines, blue for CEU, green for CHB, and red for YRI, partitioned the whole *F_PC_* space to three genealogical subspaces. **(c)** The genetic distance spectrum for Metabochip European cohorts to CEU – Northwest Europeans, FIN – Northeast European, and TSI – Southern Europeans. The nationality of the cohorts are denoted on the horizontal axis. (d) The projection for Metabochip European cohorts to the *F_PC_* space defined by CEU, FIN, and TSI reference populations. The whole space is further partitioned into three subspaces, CEU-TSI genealogical subspace (red and blue dashed lines), FIN-TSI genealogical subspace (green-blue dashed lines), and CEU-FIN genealogical subspace (red-green dashed lines), respectively. The open circles represent the mean of inferred geographic locations for the cohorts from the same country. Cohort/country codes: AF, African; AU, Australia; DE, Germany; EE, Estonia; EU, European Nations; FI, Finland; FIN, Fins in 1000 Genomes Project (1KG); FR, France; GBR, British in 1KG; GIB, Gujarati Indian in 1KG; GR Greece; Hawaii, Hawaii in USA; IBS, Iberian Population in Spain in 1KG; IT, Italy; JM, Jamaica; JPT, Japanese in 1KG; LWK, Luhya in 1KG; NO, Norway; PH, the Philippines; PK, Pakistan; SC, Seychelles; SCT, Scotland; SE, Sweden; TSI, Tuscany in 1KG; UK, United Kingdom; US, United States of America.

We next investigated whether our genetic distance method works at a much finer geographic scale. It is known that using individual-level data, principal component analysis can mirror the geographic locations for European samples^11^. Here, we analyzed the 103 GIANT European-ancestry Metabochip cohorts (48 male-only cohorts, 47 female-only cohorts, and 8 mix-sex cohorts) for fine-scale *F_st_* genetic distance measure by using the CEU, FIN, and TSI reference populations, which represent northwest, northeast, and southern European populations, respectively. For each of the GIANT European-ancestry Metabochip cohorts, *F_st_* was calculated relative to each of these three reference populations and showed concordance with the known origin of the samples (**Fig. 1c**). For example, cohorts from Finland and Estonia were close to FIN but distant to TSI; cohorts from South Europe such as Italy and Greece had small genetic distance to TSI; and cohorts from West European nations had small genetic distance to CEU. Similarly, the projected origin for each European-ancestry Metabochip cohort resembles their geographic location within the European map as expected (**Fig. 1d**). Therefore, our QC measure based upon population differentiation also works at a fine scale.

We next applied the *F_st_* genetic distance measures to 174 GIANT height GWAS cohorts (79 male-only cohorts, 76 female-only cohorts, and 19 mixed-sex cohorts; excluding Metabochip data), which were all of European ancestry imputed to the HapMap reference panel^3^. Given the three *F_st_* values to CEU, FIN, and TSI (**Fig. 2a**), the geographic origin for each cohort can be inferred as for the GIANT BMI Metabochip data (**Supplementary notes**). The projected coordinates of each GWAS cohort matches its origin very well (**Fig. 2b**). For example, a Canadian cohort, the Quebec Family Study (QFS), was closely located to DESIR, a French cohort, consistent with the French genetic heritage of the QFS^16^. In addition, we also observe complexity due to mixed samples from different countries. For example, the DGI/Botnia study had samples recruited from Sweden and Finland, and its inferred geographic location is in between of the Swedish cohorts and Finnish cohorts^17^. We also note that for the MIGEN consortia cohorts, which are from Finland, Sweden, Spain and the US, the same allele frequencies were reported for all their sub-cohorts, and all cohorts were allocated to southern Europe (very closely located to 1KG IBS cohort; **Fig. 2b and Supplementary Fig. 2**). As the allele frequencies, used in QC steps to eliminate low quality loci, were not directly used in estimating genetic effects in the GWAMA, the reported allele frequencies in MIGEN have not impacted on the published GWAMA results^3^.

**Figure 2.**
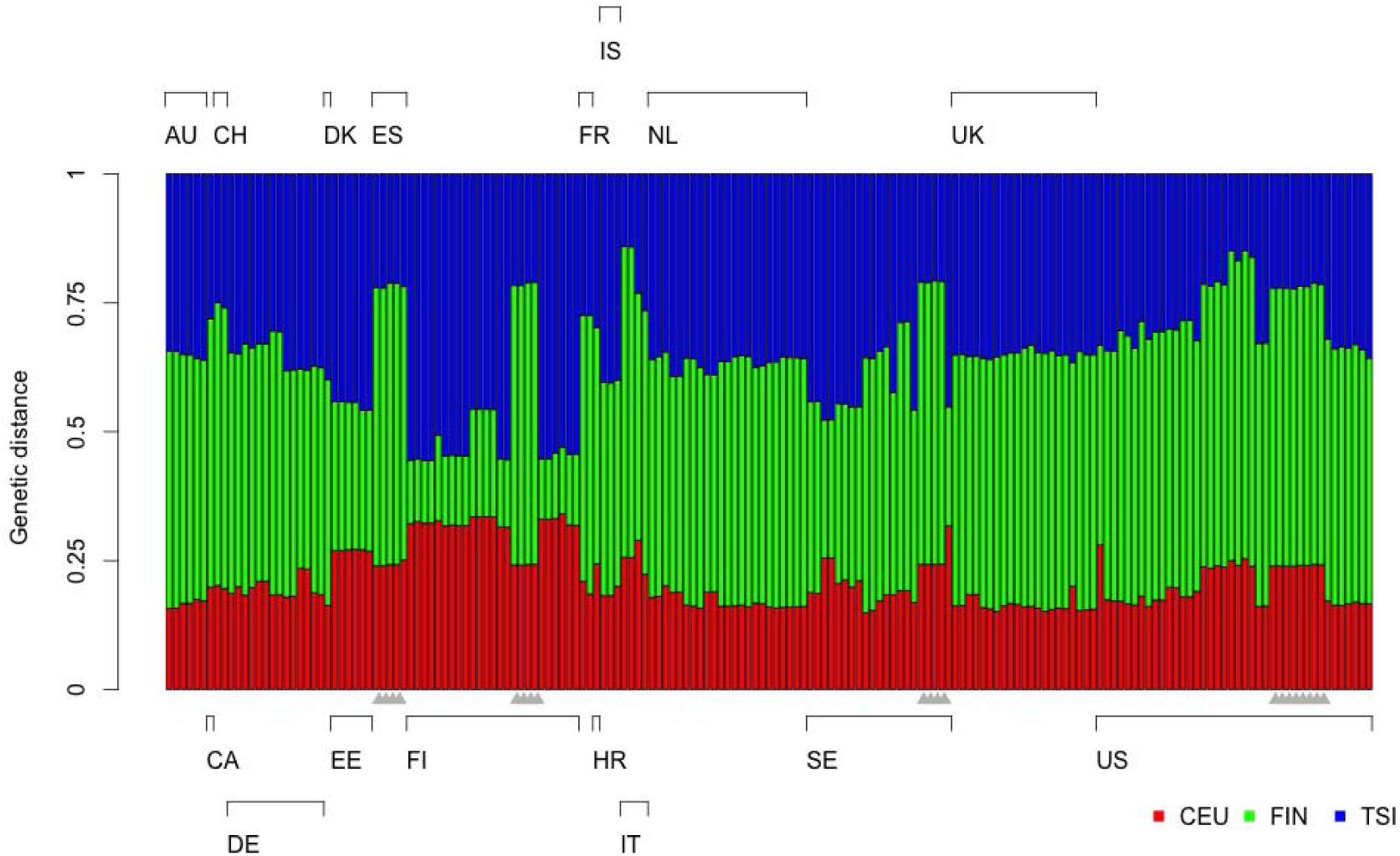
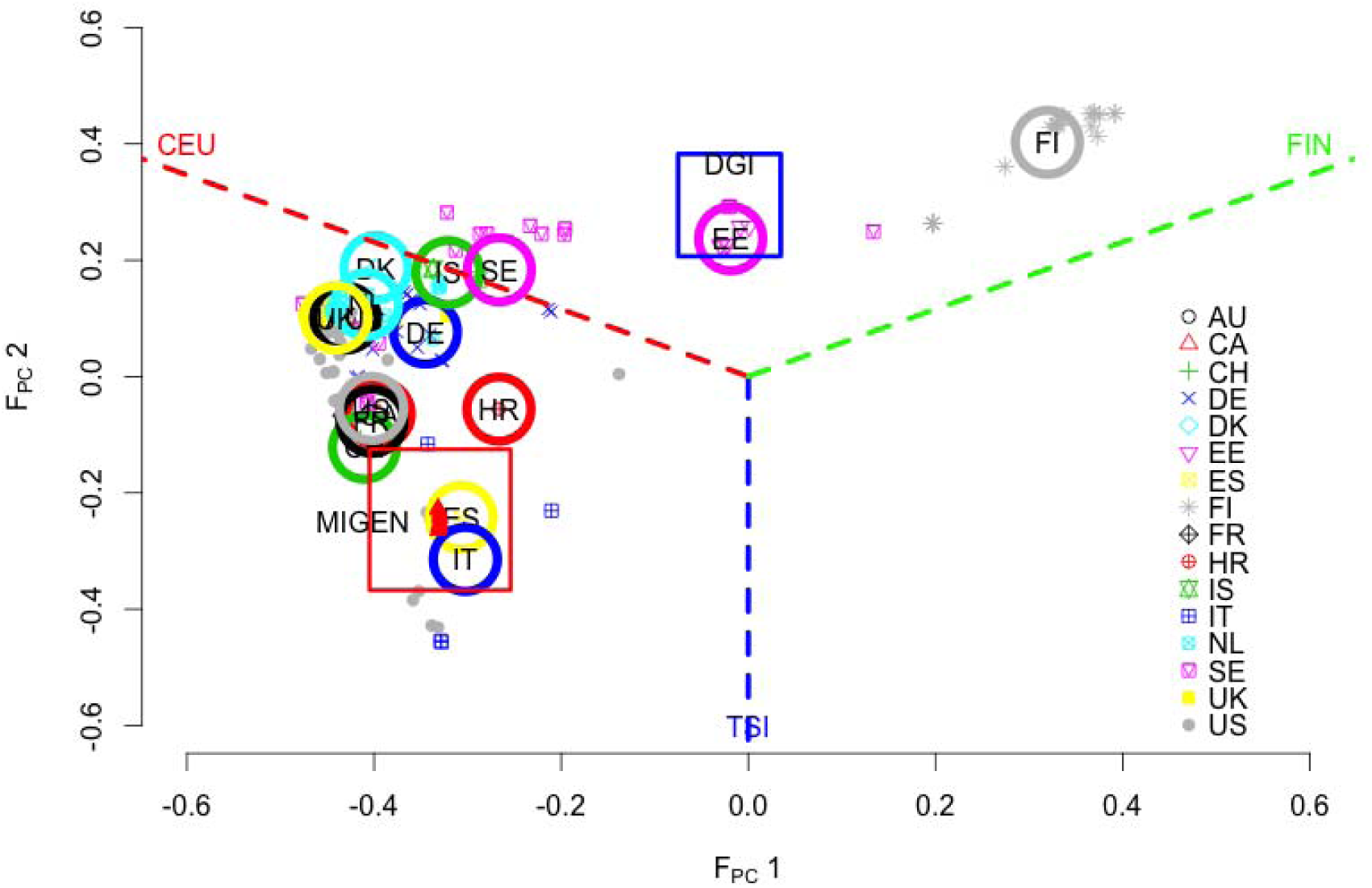
Using the genetic distance spectrum to infer the geographic origins for GIANT height GWAS cohorts. **(a)** Each cohort has three *F_st_* values by comparing with CEU, FIN, and TSI reference samples. The height of each bar represents its relative genetic distance to these three reference populations. The nationalities of the cohorts were denoted along the horizontal axis. The grey triangles along the x-axis indicate MIGEN cohorts. **(b)** Given the three *F_st_* values, the location of each cohort can be mapped. The whole space was partitioned into three subspaces, CEU-TSI genealogical subspace (red and blue dashed lines), FIN-TSI genealogical subspace (green and blue dashed lines), and CEU-FIN genealogical subspace (red and green dashed lines). DGI (in the blue box) had samples from the Botnia study. Across MIGEN cohorts (denoted as red triangles in the red box), the same allele frequencies (likely calculated from a South European cohort) were presented for each cohort. Cohort/country codes: AU, Australia; CA, Canada; CH, Switzerland; DE, Germany; DK, Denmark; EE, Estonia; ES, Iberian Population in Spain in 1KG; FI, Finland; FR, France; GR, Greece; IT, Italy; IS, Iceland; NL, Netherlands; SE, Sweden; UK, United Kingdom; US, United States of America.

Next, we show that *F_st_* can detect populations that have a different demographic past. Using all 1KG European samples as the reference panel (that is, an “averaged” European reference panel), most cohorts in GIANT had < 0.005 with this average, which agrees with previously reported results using individual level data from European nations^11^. A few cohorts showed large such as the AMISH cohort with *F_st_ =* 0.018, and the North Swedish Population Health Study (NSPHS)^18^ with *F_st_ =* 0.014. Consistent with these results, both these populations are known to have been genetically isolated (**Supplementary Fig. 3**).

### Principal component analysis for allele frequencies

It is well established that given individual-level data principal component analysis (PCA) can reveal the ancestral information for samples^12^. Given the same allele frequencies as used for *F_st_*-based analysis above, we conducted PCA for allele frequencies, denoted as meta-PCA. In meta-PCA each cohort was analogously considered as an “individual”. For example, 120 Metabochip cohorts were considered as a sample of 120 “individuals”. Although the inferred ancestral information was for each cohort rather than any individuals, implementation of meta-PCA was the same as the conventional PCA (**Supplementary Notes**).

Meta-PCA was tested with 1KG samples over nearly 1 million SNPs. The cohort-level allele frequencies were calculated first for 26 1KG cohorts, and meta-PCA was conducted. The projected cohorts were consistent to their genetic origin (**Fig. 3**). In contrast, conventional PCA was also conducted on 1KG individual genotypes directly, and the mean coordinates for each cohort was then calculated. As illustrated in Fig. 3, these two techniques resulted in nearly identical projection for 1KG, and the correlation between cohort coordinates remained consistently high for the first eight eigenvectors, *R^2^ >* 0.8. It indicated that meta-PCA could reveal genetic background for each cohort as precise as that based on individual-level data.

**Figure 3.**
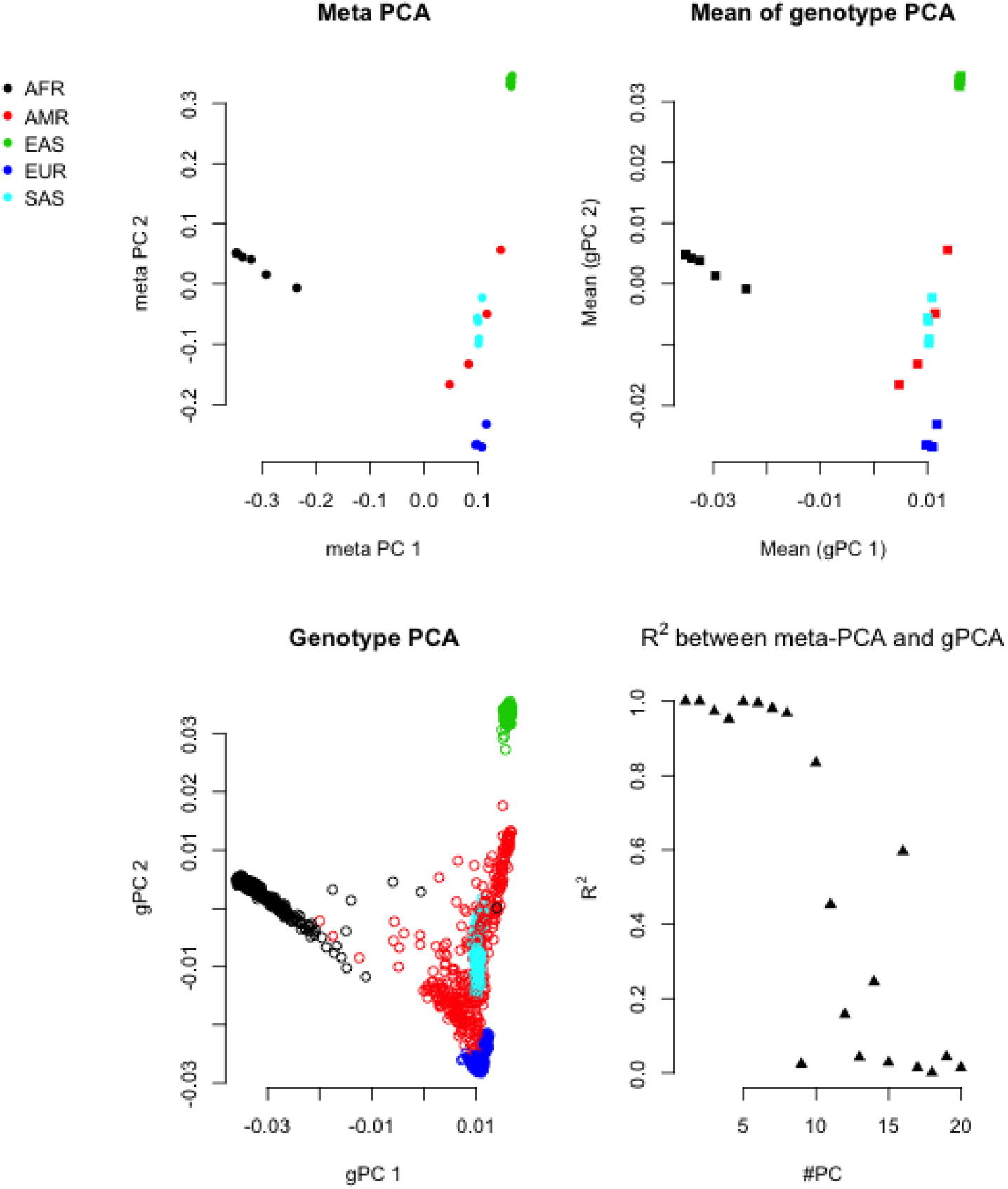
Comparison between Meta-PCA and genotype PCA on 1KG. Top left panel is the projection of cohorts based on cohort-level allele frequency for 1KG samples on the first two eigenvectors. Bottom left panel is conventional PCA based on individual genotypes on the first two eigenvectors. Top right panel is the projection by taking the mean of the 1KG individuals within each cohort. Bottom right panel is the correlation, measured in, between meta-PCA and genotype PCA for the first twenty eigenvectors.

We applied meta-PCA to 120 Metabochip cohorts for nearly 34 thousand common SNPs between Metabochip and 1KG variants, with the inclusion of 10 1KG cohorts (East Asian: CHB, JPT; South Asian: GIH; European: CEU, FIN, GBR, IBS, TSI; African: LWK, YRI) as the reference cohorts. Consistent with demographic information, the inferred ancestral information of each cohort agreed well with demographic information. For example, PROMISE (Pakistan) located very close to GIH, CLHNS (Philippines) close to CHB and JPT, ARIC (African American) and SPT (Jamaican) close to YRI and LWK, and the European cohorts close to CEU and FIN (**Fig 4**).

**Figure 4.**
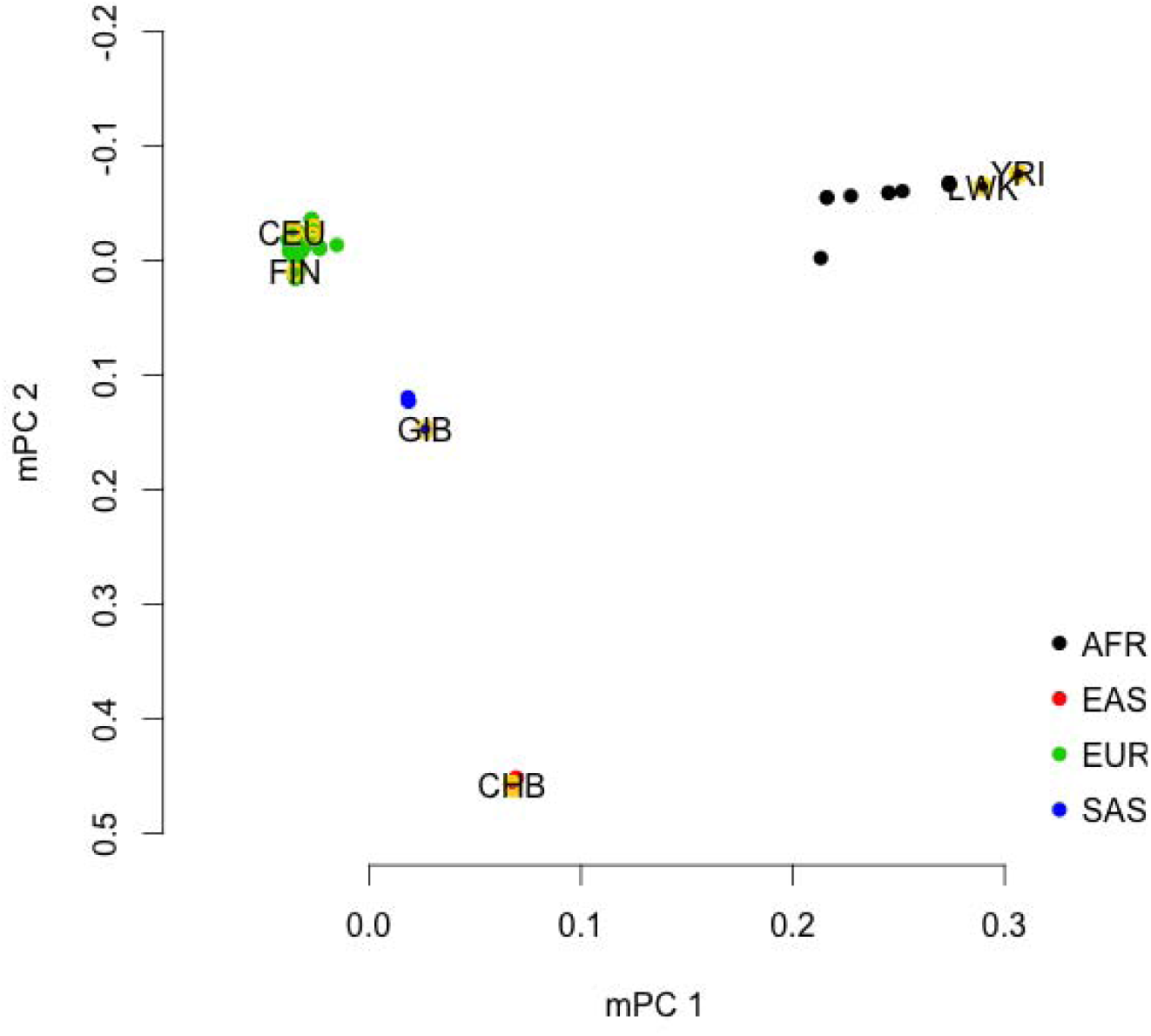
Recovery of cohort-level genetic background for GIANT BMI Metabochip cohorts using meta-PCA. The x-axis and y-axis represent the first two eigenvectors from meta-PCA. In meta-PCA, Metabochip cohorts could be classified into African ancestry (AFR), European ancestry (EAS), East Asian Ancestry (EAS), and South Asian Ancestry (SAS). The 1KG cohorts, yellow open circles, were added for comparison.

We also applied meta-PCA to 174 GIANT height GWAS cohorts for nearly 1M SNPs, with the inclusion of 10 1KG reference cohorts. At the global-population level, the 174 cohorts were all allocated close to CEU and FIN, consistent with their reported demographic information (**Fig. 5**). For fine-scale inference, we conducted meta-PCA again but with the inclusion of the five European samples. As demonstrated, the resolution of the inferred relative location between European cohorts reflected their real geographical locations, as previously observed using individual-level data^11^.

**Figure 5.**
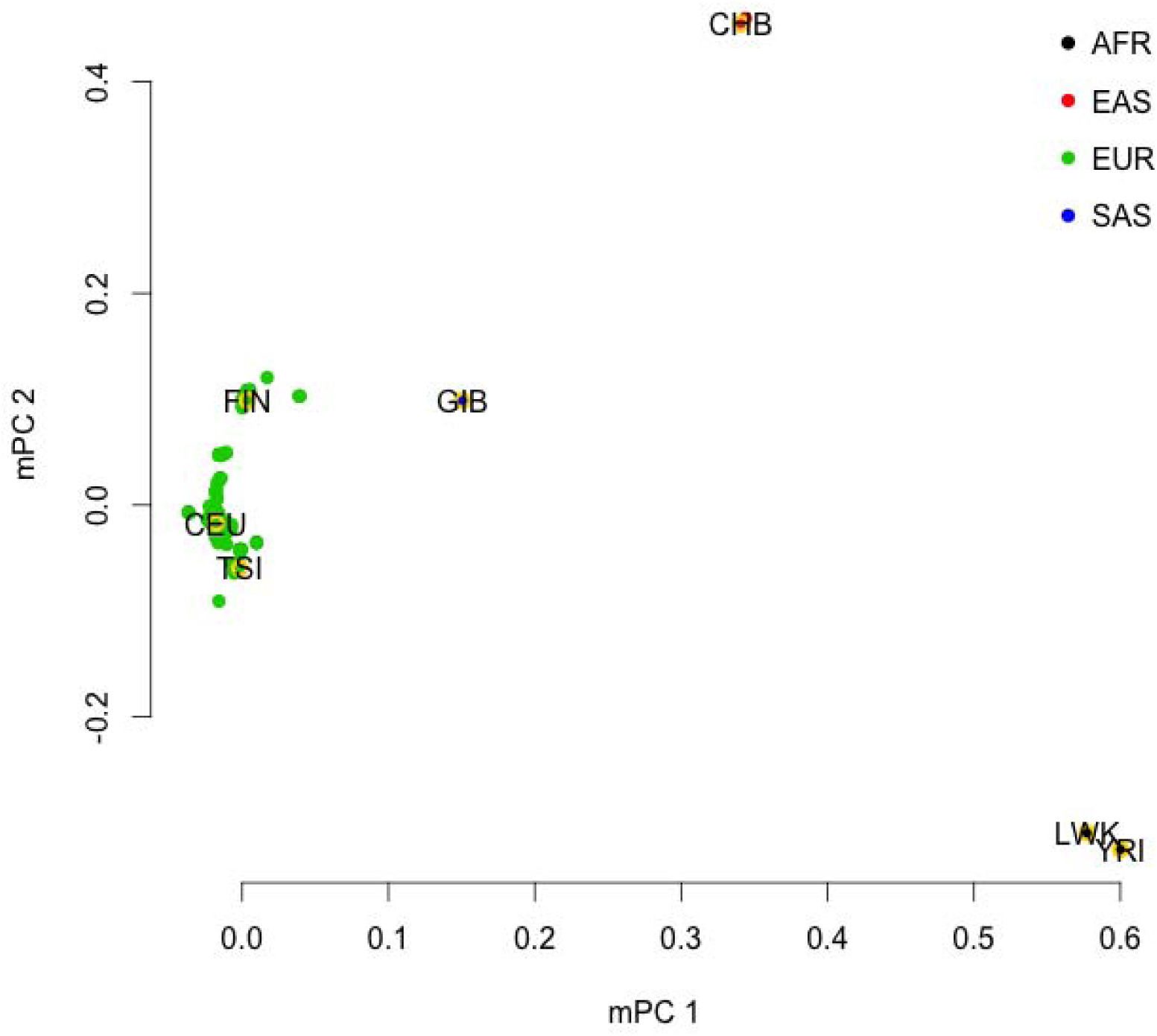
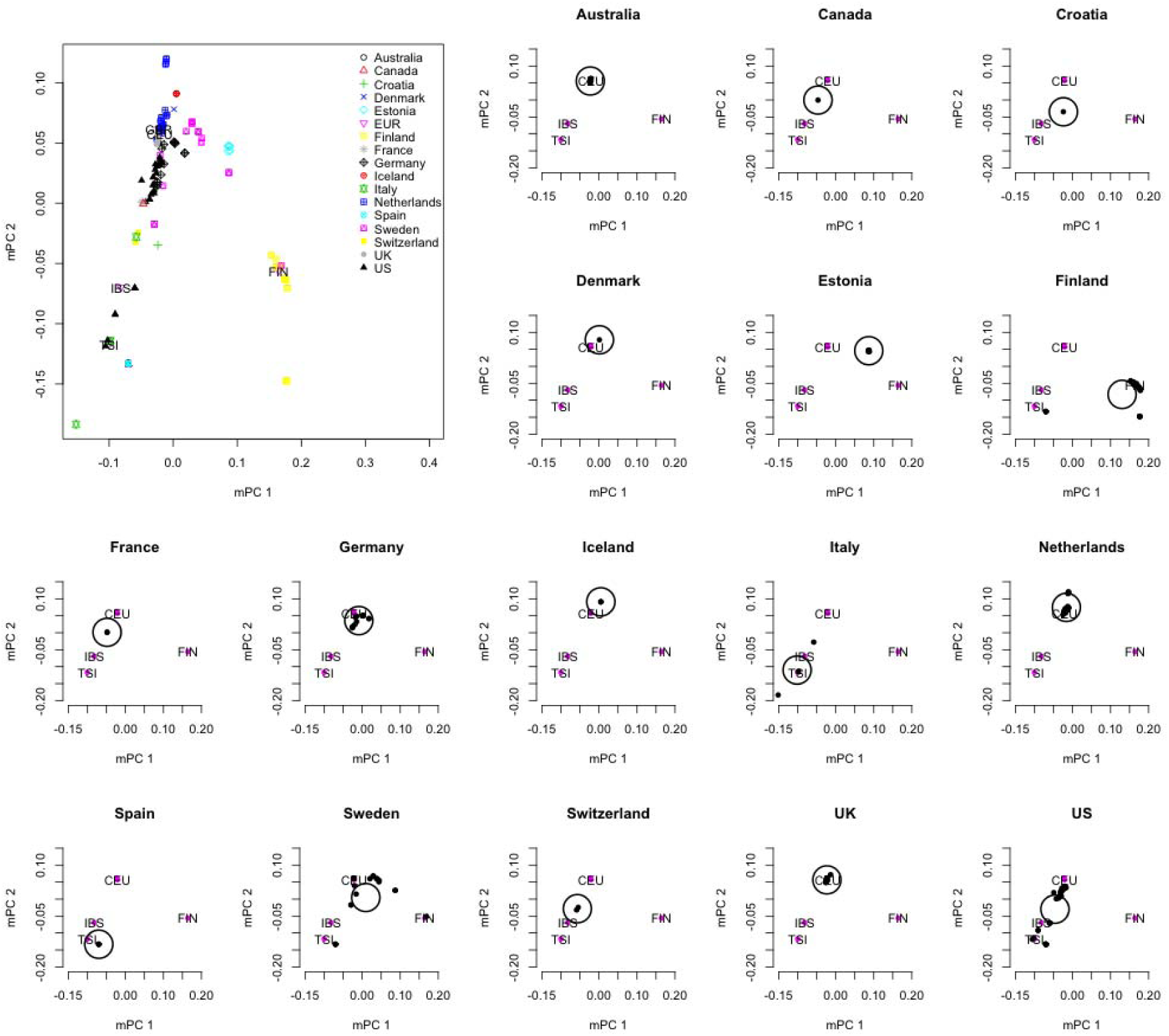
The recovery of cohort-level genetic background using meta-PCA analysis for GWAS height cohorts. The x-axis and y-axis represent the firs two eigenvectors inferred from meta-PCA. a) The genetic background inferred with the inclusion of 10 1KG reference populations. b) The genetic background and relative geographic location for 174 GIANT height cohorts. The large plot on top left was an overview of 174 cohorts, and the rest of plots were classified by the reported demographic information of cohorts. Within each country-level plot, the small black points represent one cohort, and the large open circle the mean coordinates for those cohorts from the same country.

These results were consistent to what observed from *F_pc_* as described in the last section, and also agreed well with demographic information. So, based on the reported allele frequencies, the demographic information could be examined by meta-PCA method.

### λ*_meta_* to detect pairwise cohort heterogeneity and sample overlap

For a single cohort GWAS, *λ_GC_* provides a tool for assessing average trait-SNP associations in GWAS^9^, and an value departing from 1 may indicate undesired phenomena such as population stratification. In this study, we use the summary statistics for a pair of cohorts to calculate *λ_meta_*, a metric that examines heterogeneity from the concordance of reported effect sizes and sampling variance. We use 30,000 markers in linkage equilibrium along the genome between a pair of cohorts to estimate *λ_meta_*.

For a SNP marker (*i*), given its reported estimated effect size (*b_i_*) and sampling variance 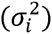 in a pair of cohorts 1 and 2, we can calculate a test statistic 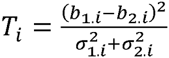, the ratio between the squared difference of their reported effects to the sum of their reported sampling variances. Under the null hypothesis of no overlapping samples/heterogeneity, *T* follows a chi-square distribution with 1 degree of freedom

(**Supplementary notes**). 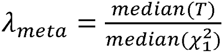, the ratio between the median of the 30,000 *T* values and the median of a chi-square statistic with 1 degree of freedom (a value of 0.455), has an expected value of 1 for two independent GWAS summary statistics sets for the same trait. When there is heterogeneity between estimated genetic effects, the expectation is *λ_meta_* > 1, and in contrast *λ_meta_* <1 if there are overlapping samples. In general, not only overlapping samples but also close relatives present in different cohorts can lead to correlated summary statistics generating *λ_meta_* <1 (**Supplementary notes**). However, unless the proportion of overlapping relatives is substantial and their phenotypic correlation is high, the correlation of the summary statistics due to the effective number of overlapping samples (*n_o_*) is expected to be dominated by the same individuals contributing phenotypic and genetic information to different cohorts (**Supplementary Fig. 4**). Furthermore, if genomic control is applied to adjust the sampling variance^19^ then *λ_meta_* will be reduced relative to its value without genomic control (**Supplementary notes**).

We estimated *λ_meta_* from published GWAS summary statistics for a range of traits (other than BMI and height) and were able to find examples of both deflated and inflated *λ_meta_*. First, we tested the *λ_meta_* on data sets with known overlap. For example, GWAS summary statistics for schizophrenia were available in two phases: the first had 9,394 controls and 12,462 cases^20^, and in the next phase about 18,000 Swedish samples were added^21^. Such a substantial overlap sample between these two sets of summary statistics led to the estimated value of *λ_meta_* as low as 0.257 (**Supplementary Fig. 5**), consistent with this known overlap. In contrast, heterogeneity between data sets (represented by *λ_meta_* > 1), was observed between GWAS summary statistics of rheumatoid arthritis from European and Asian studies^22^, for which *λ_meta_* = 1.09 (**Supplementary Fig. 6**). In addition, we note that the distribution of the empirical *T*-statistics deviates from expectation at the upper tail of the distribution, suggesting differences in effect size or linkage disequilibrium between these two ancestries.

Next, we estimated *λ_meta_* from pairs of cohorts from the 174 GIANT height GWAS cohort^3^. We found no evidence for substantial sample overlap but do observe between-cohort heterogeneity, and technical artifacts. From the 174 GIANT height GWAS (supplied data files)^3^, we calculated 15,051 cohort-pairwise *λ_meta_* values, resulting in a bell-shape distribution (**Fig. 6a,b**) with the mean of 1.013 and the empirical standard deviation (S.D.) of 0.022, which was greater than theoretical S.D. of 0.014. The empirical mean and S.D can be used to construct a z-score test for each *λ_meta_*. These results are consistent with a small amount of heterogeneity, which is not unexpected due to variation of actual (unknown) genetic architecture and analysis protocols. However, the mean is close to 1.0 and based upon this QC metric the results are consistent with stringent quality control and data cleaning. The minimum *λ_meta_* value was around 0.88 (between SORBS MEN and SORBS WOMEN, **Fig. 3c**), with *p*-value < 1e-10 (testing for the difference from 1), and the maximum was 1.245 (between SardiNIA and WGHS, **Fig. 6d**), with *p*-value < 1e-10, leading to the most deflated and inflated across GIANT height study cohorts; both were significant after correction for multiple testing. Illustrating *λ_meta_* (**Fig. 6b**) highlighted that 20 cohorts from the MIGEN consortium showed substantially lower *λ_meta_* with many other cohorts (right-bottom triangle in **Fig. 6b**) than the average, consistent with over-conservative models for statistical association analyses being used in these cohorts – which may be due to very small sample size (ranging from 36 to 320 for the 20 MIGEN cohorts, with an average sample size of 132). Consistent with this, cohorts from MIGEN also have many of their *λ_gc_* < 1 (**Fig. 7a**). In contrast, the SardiNIA cohort (4,303 samples) showed heterogeneity with nearly all other cohorts (**Fig. 7b**), perhaps due to unknown artifacts or a slightly different genetic architecture for height as result of demographic history^23^.

**Figure 6.**
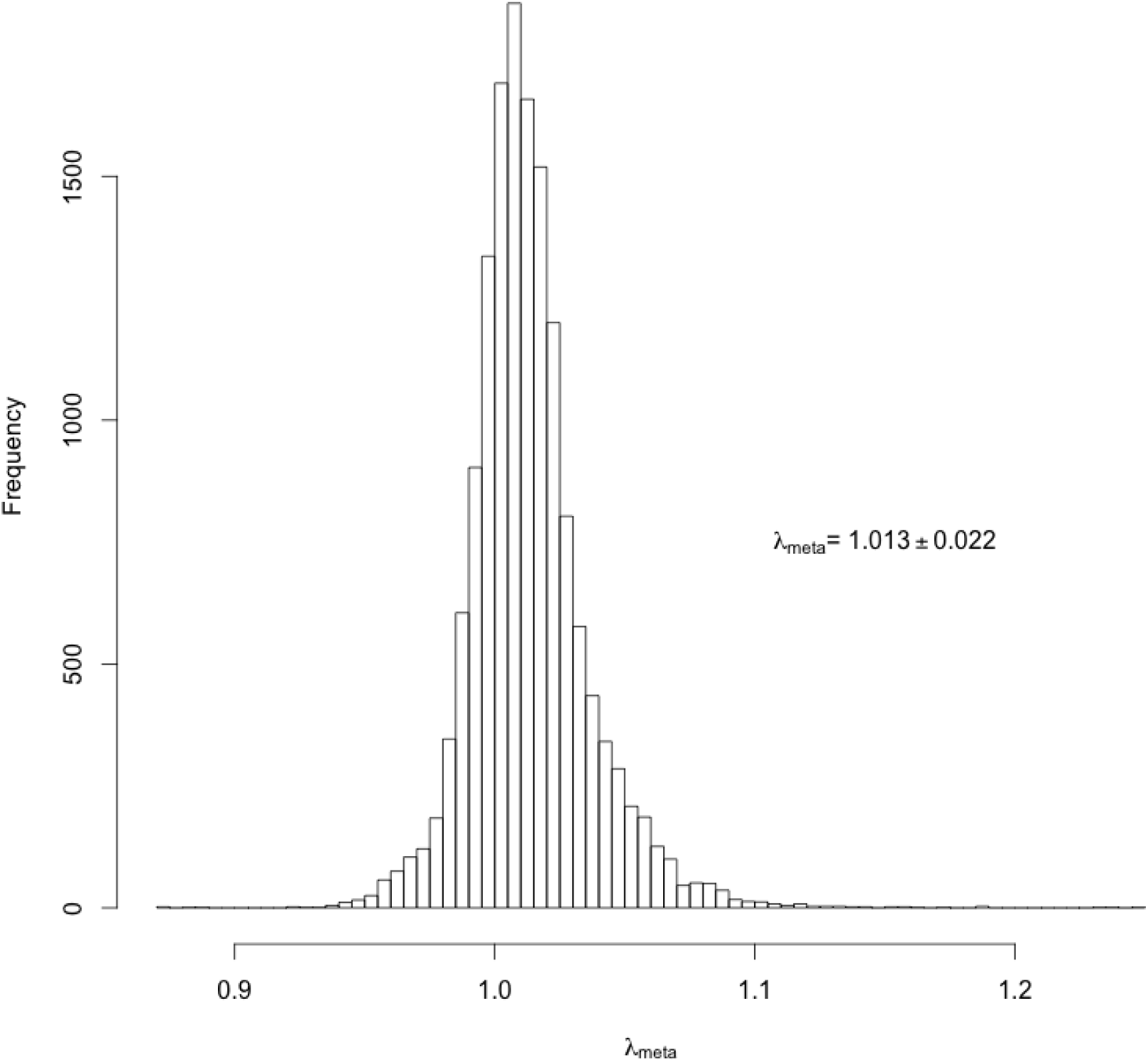
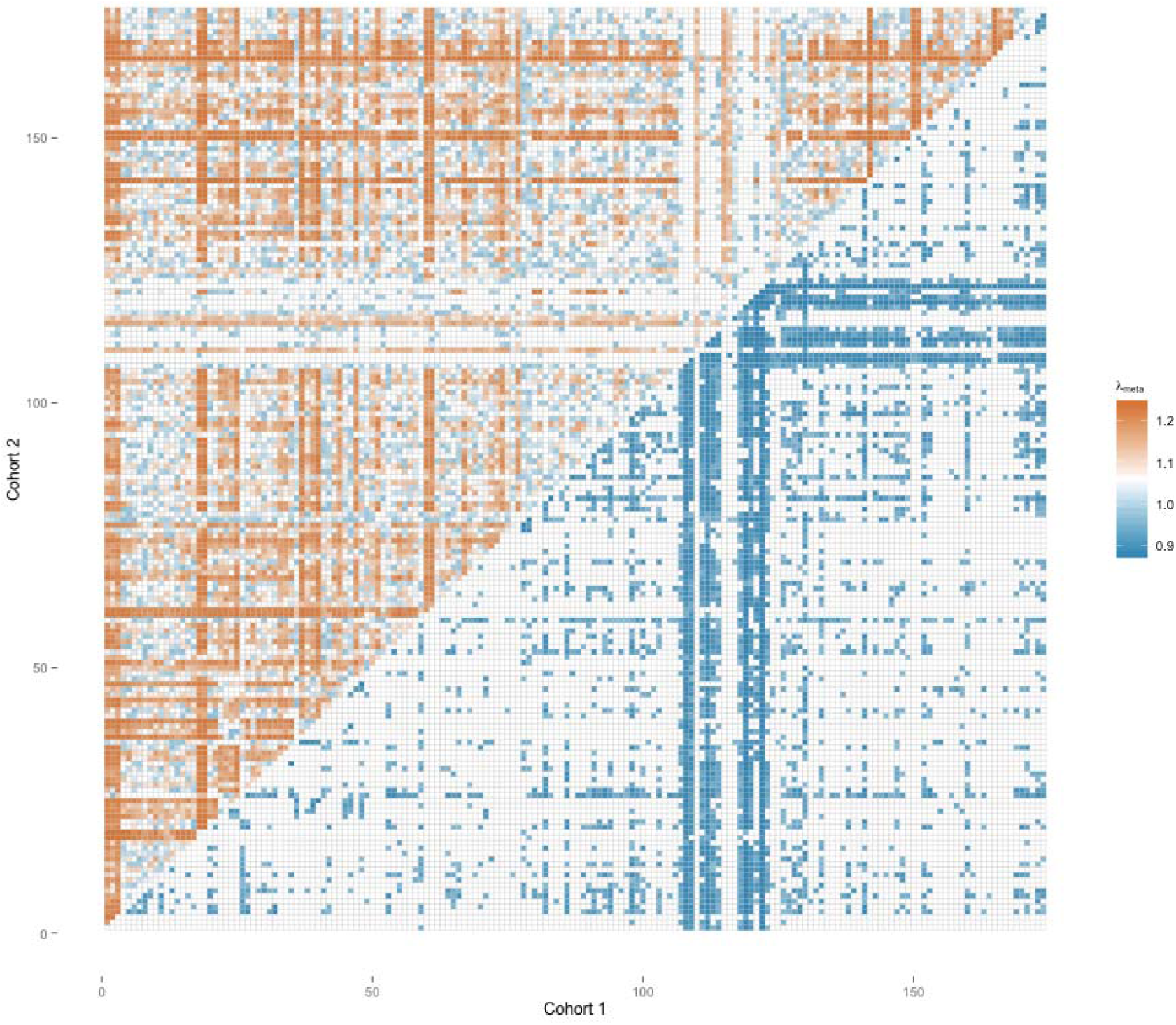
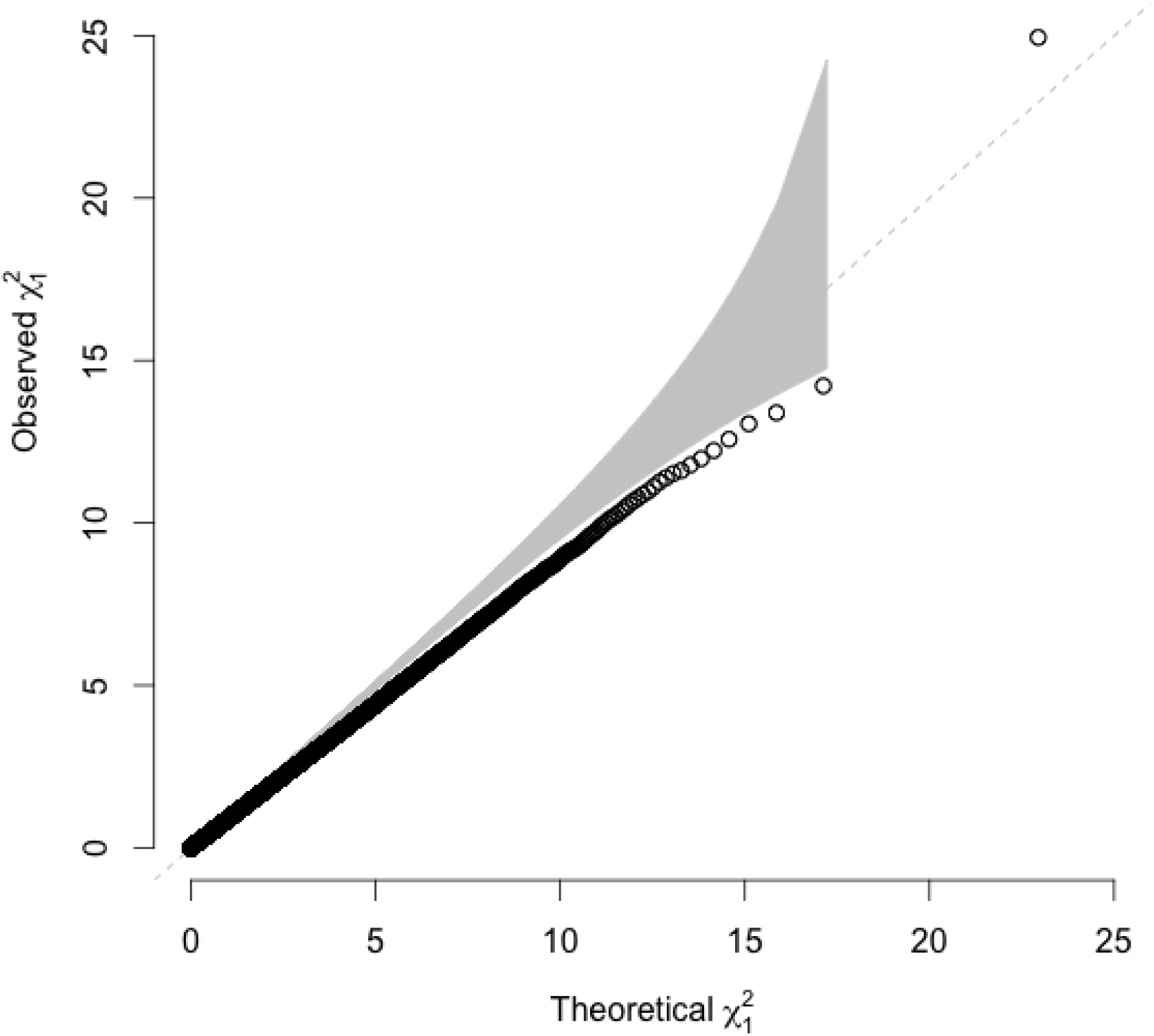
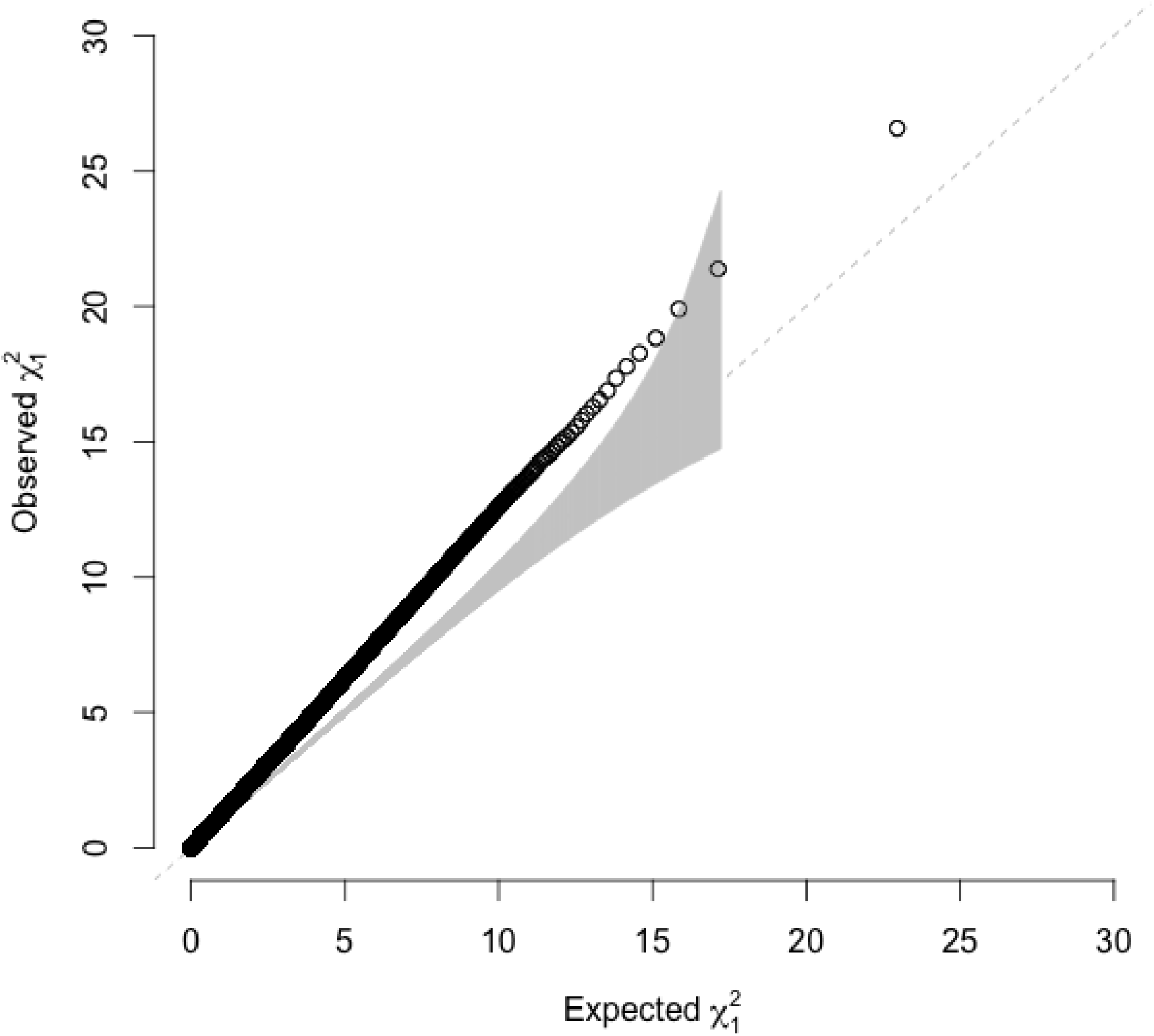
*λ_meta_* for the GIANT height GWAS cohorts. Given 174 cohorts, there are 15,051 *λ_meta_* values, which provide the overview of the quality control of the summary statistics. **(a)** The distribution of *λ_meta_* from 174 cohorts/files used in the GIANT height meta-analysis. The overall mean of 15,051 *λ_meta_* is 1.013, and standard deviation is 0.022. **(b)** The heat map for *λ_meta_*. Cohorts showed heterogeneity (*λ_meta_* > 1) are illustrated on left-top triangle, and homogeneity (*λ_meta_* < 1) on right-bottom triangle. **(c)** Illustration for homogeneity between two cohorts (SORBS MEN & WOMEN), *λ_meta_ <* 0.876. **(d)** Illustration of SARDINIA & WGHS, this pair of cohorts has *λ_meta_ =* 1.245. The grey band represents 95% confidence interval for *λ_meta_*.

**Figure 7.**
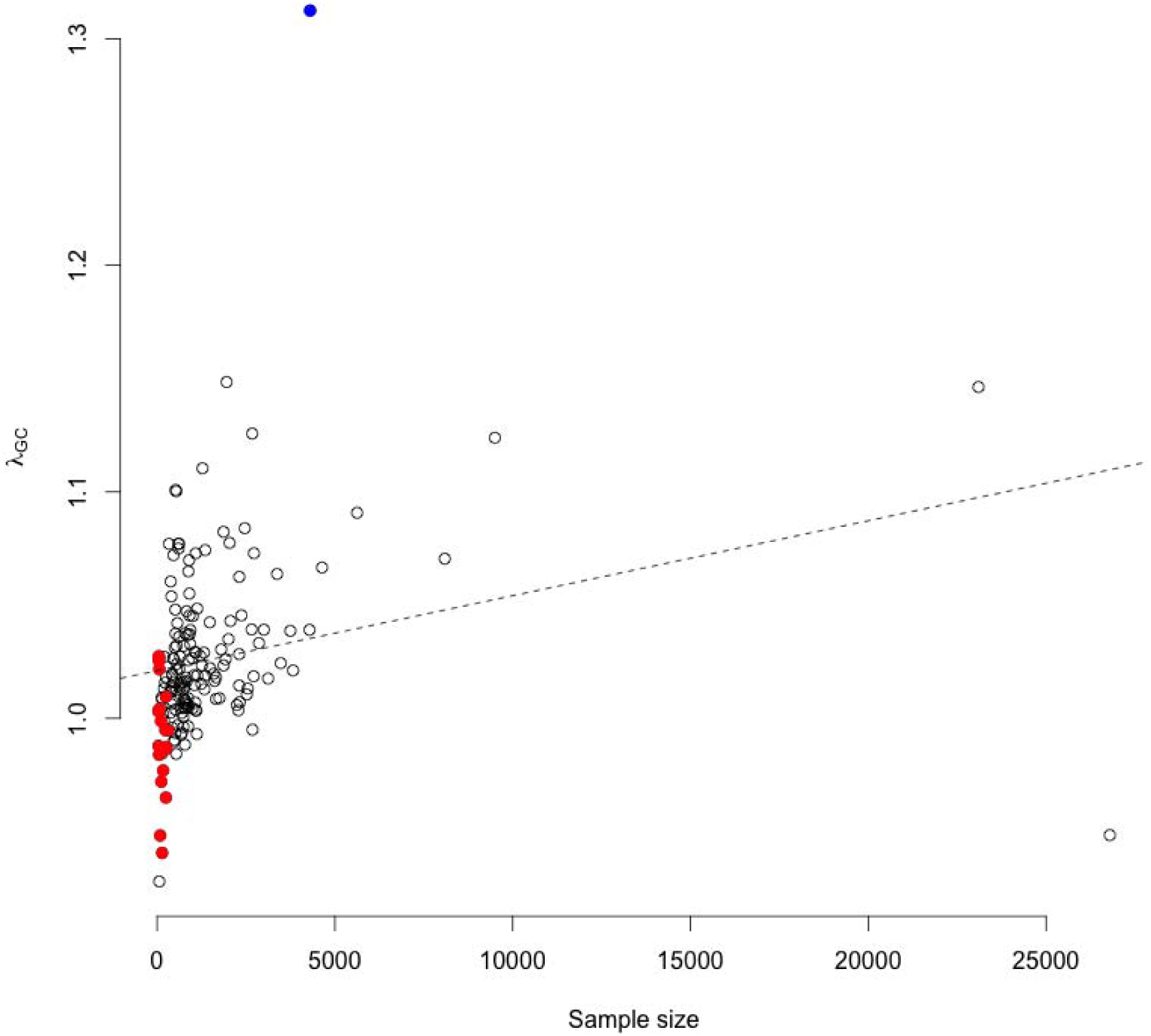
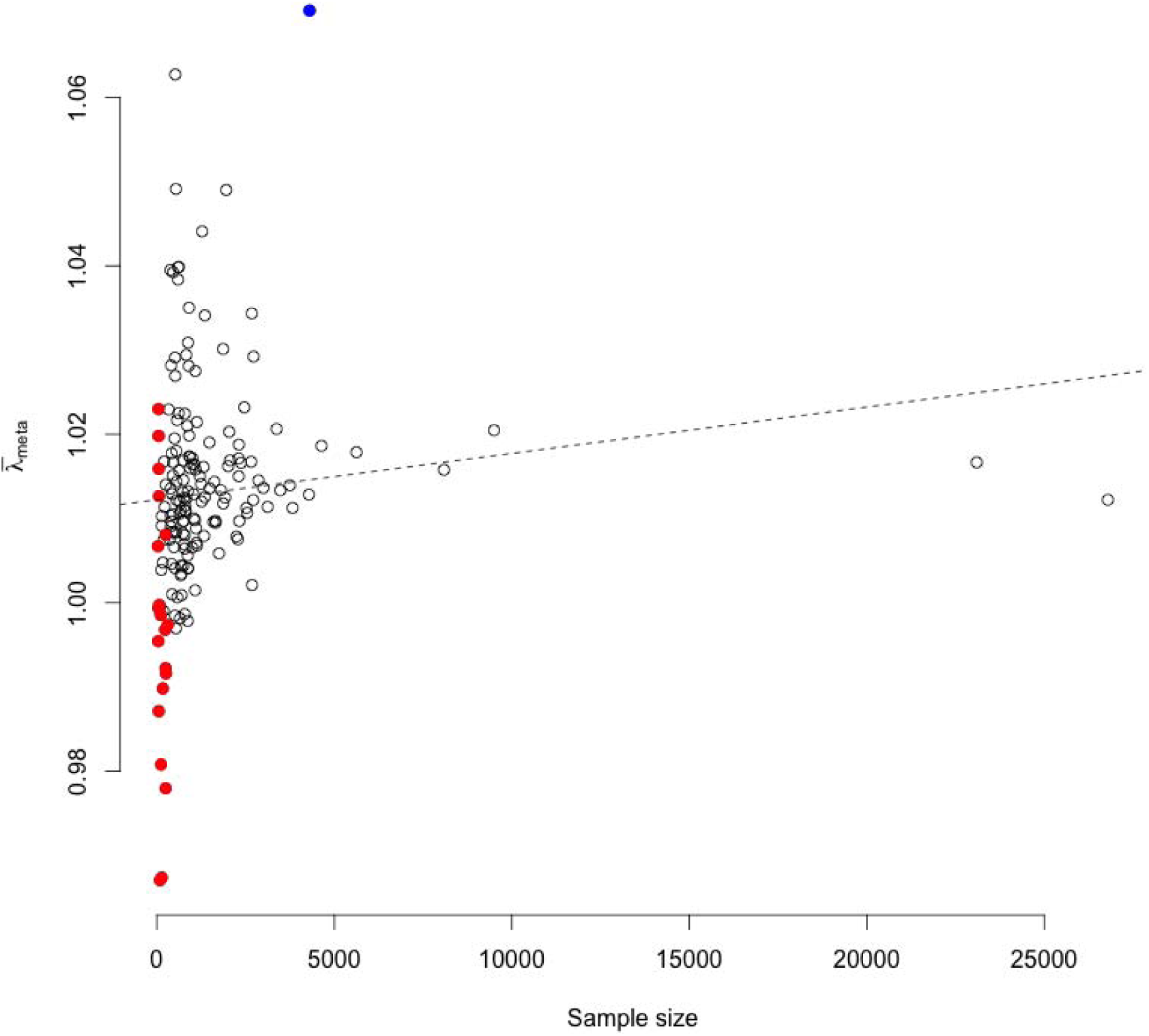
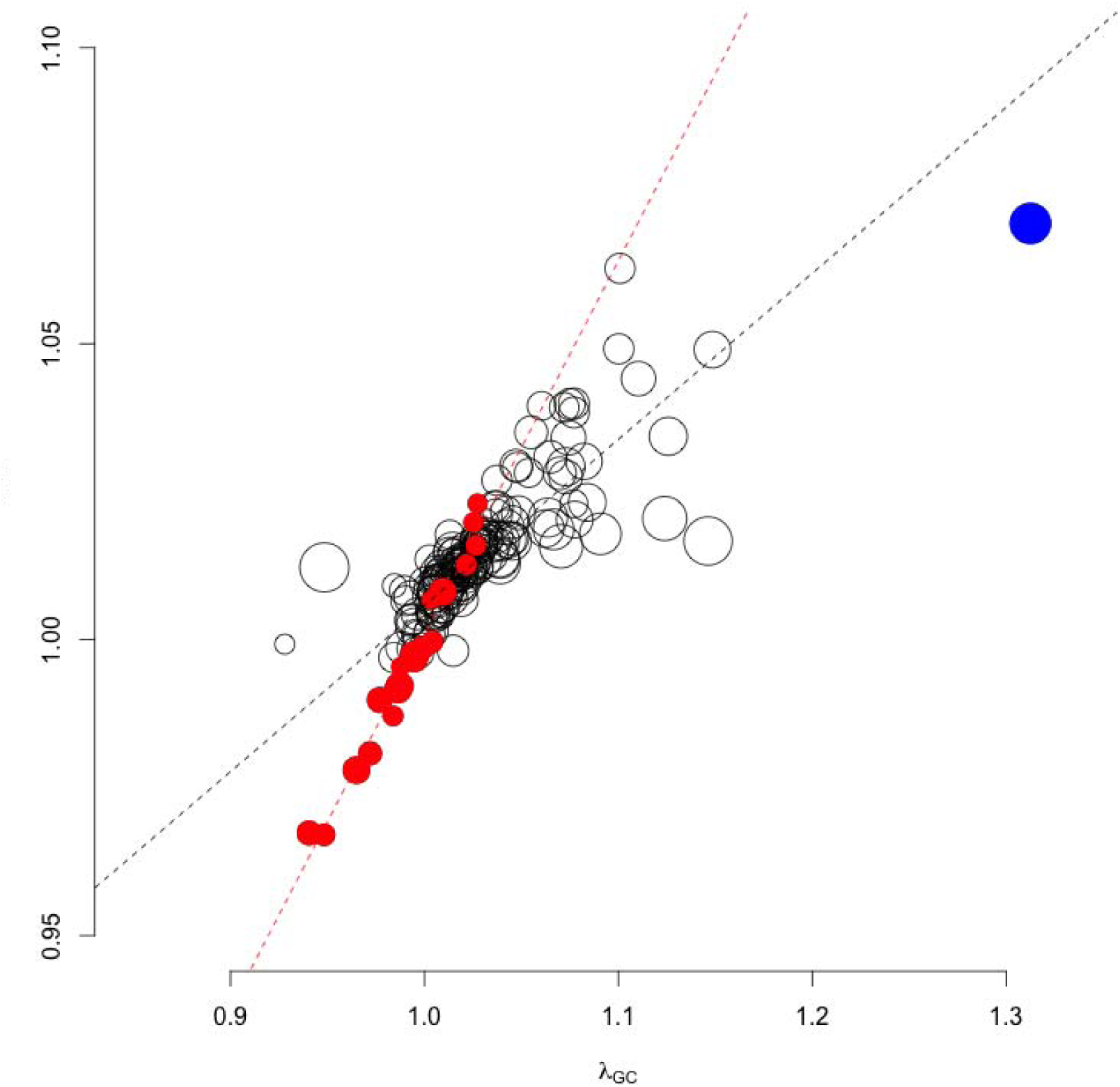
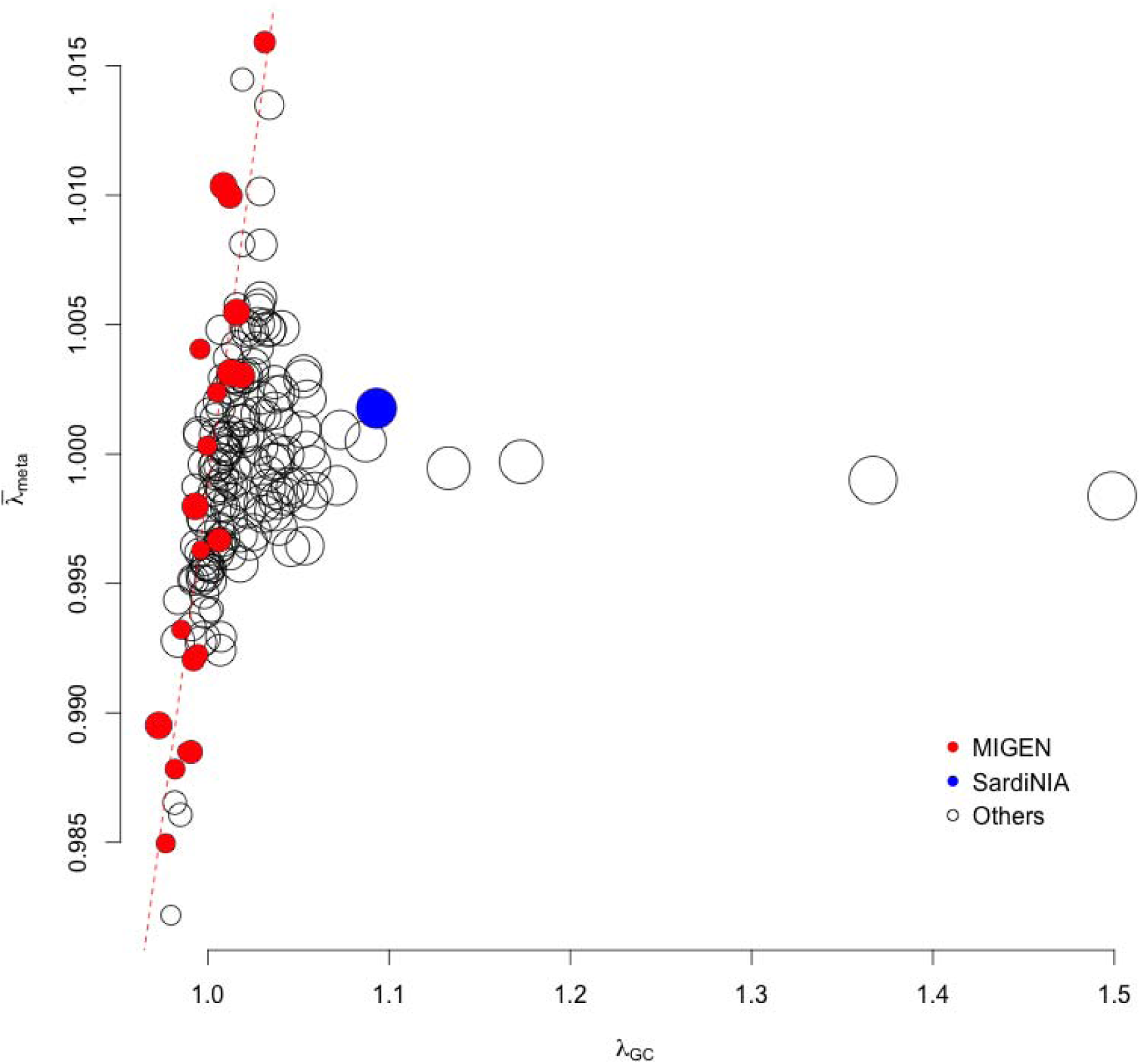
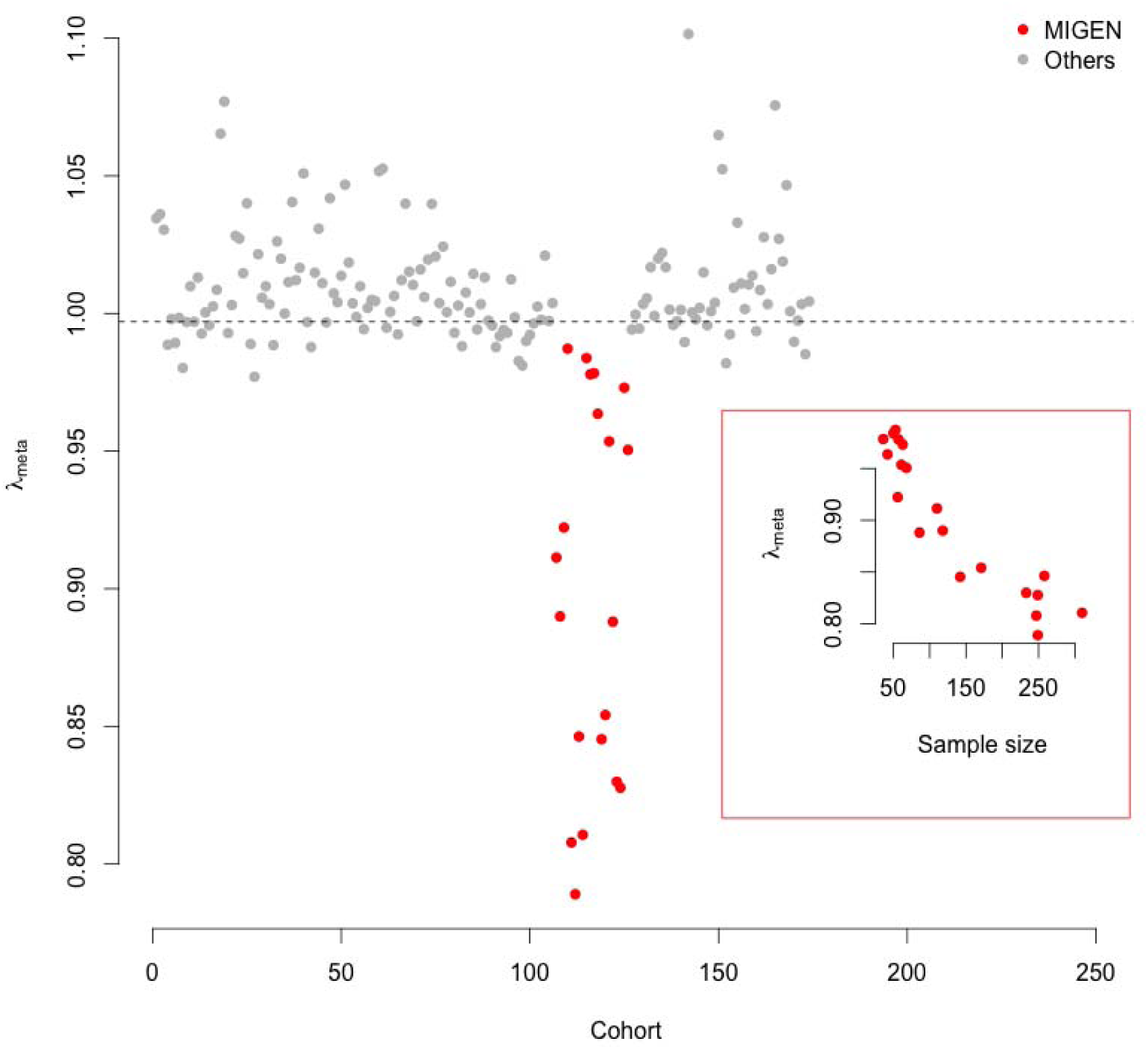
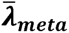 and *λ_gc_* for GIANT height GWAS cohorts. **(a)** Sample size of each cohort against *λ_GC_*. The linear regression is presented as a dashed line, *λ_GC_* = 1.021 + 0.0000033N, and *R^2^ =* 0.013. **(b)** Sample size of each cohort against 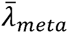, which was the mean of a cohort’s *λ_meta_* over all other cohorts. The linear regression is presented as a dashed line, 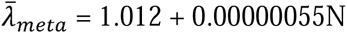 (N = reported sample size), and *R^2^ =* 0.055. **(c)** *λ_GC_* against 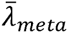 for each cohort, showing a strong correlation, *R^2^ =* 0.70. The black dash line indicates the regression slope for all 174 pairs: 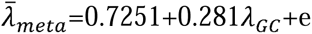. The red dashed line indicates the regression slope for 20 pairs of MIGEN cohorts: 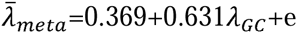. The side of each circle is proportional to sampling size on logarithm scale. **(d)** Small sample size leads to a correlation between 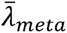 and *λ_GC_* using 174 GIANT height GWAS sample size. 30,000 independent loci, minor allele frequency ranged from 0.1~0.5, were simulated, and *h^2^ =* 0.5. The red dashed line indicates the regression slope for 20 simulated MIGEN cohorts, 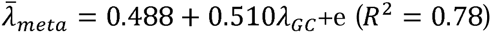. The side of each circle is proportional to sampling size on logarithm scale. **(e)** *λ_meta_* for whole MIGEN to 174 cohorts. 20 MIGEN files were combined together to make “whole MIGEN” via meta-analysis, and the summary statistics were used to calculate *λ_meta_* with 174 cohorts using 30,000 independent loci. As MIGEN cohorts were part of “whole MIGEN”, their *λ_meta_* were in general below 1. The dashed line is the mean of *λ_meta_* of the “whole MIGEN”. The subplot (red box) shows a strong correlation of 0.93 between *λ_meta_* (for “whole MIGEN” vs each MIGEN cohort), and sample size of each MIGEN cohort.

We investigated the relationship between 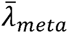 (the mean of all *λ_meta_* values of a given cohort with each of the other 173 GIANT height cohorts) and *λ_GC_* among the GIANT height cohorts. If there are no technical issues, such as inflated or deflated sampling variance for the estimated effects, we would expect to see: i) a correlation between *λ_GC_* and sample size; ii) no correlation between 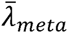 and sample size; iii) no correlation between 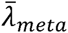 and *λ_GC_* (**Supplementary Fig. S7**). Consistent with a previous study^24^, for a polygenic trait such as height *λ_GC_* of each cohort was related to its sample size (correlation of 0.235, *p* = 0.0018). In contrast, the correlation between *λ_meta_* and sample size was of 0.116 (*p* = 0.127) (**Fig 7a,b**). Nevertheless, the correlation between the mean of 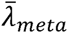 and *λ_GC_* was 0.836 (*p*<10e-16) for 174 GIANT height cohorts (**Fig 7c**). We note that the 20 MIGEN cohorts had proportionally small *λ_GC_* and 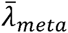, with very high correlation between them (*ρ =* 0.98); in contrast, the SardiNIA cohort, which had the largest *λ_GC_*, showed the largest 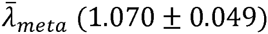, standing out as a special case among the GIANT height cohorts. Assuming a polygenic model of *h^2^ =* 0.5 over 30,000 independent loci, we simulated 174 cohorts using the actual size samples from the GIANT height cohorts (**Supplementary notes**), and observed an increased correlation (*R^2^* = 0.78) between 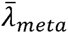 and *λ_GC_* for simulated cohorts with sample sizes of the MIGEN cohorts **(Fig. 7d)**. Other effects, such as inflated/deflated sampling variance of the estimated genetic effects could also lead to correlation between 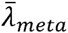 and *λ_GC_* (**Supplementary Fig. S8**). In addition, we constructed a single MIGEN analysis by combining the 20 MIGEN cohorts using an inverse variance weighted meta-analysis^25^, and calculated *λ_meta_* between this combined MIGEN cohort and all 174 cohorts. As expected, the combined MIGEN had *λ_meta_* = 0.90 ± 0.07 with 20 MIGEN cohorts due to overlapping samples. In contrast, *λ_meta_ =* 1.01 ± 0.02 with 154 other cohorts, was consistent with neither heterogeneity nor sample overlap. Given that the MIGEN (2,340 samples) and SardiNIA (4,303 samples) cohorts contributed less than 3% of the total sample size (253,288 samples from the GIANT height GWAS cohorts), any impact of unusual *λ_meta_* values on the meta-analysis results is very small. Given no heterogeneity between a pair of cohorts, a deflated *λ_meta_* reflects the effective number of overlapping samples (**Supplementary notes**). For example, the “combined MIGEN” had *λ_meta_* values proportional to the sample size of each MIGEN cohort (**Fig 7e**).

The statistical power of detection of overlapping samples is maximized when a pair of cohorts has equal sample size (**Fig. 8a**), or in other words the confidence interval for null hypothesis of no overlapping samples depends on the sample sizes for a pair of cohorts. As a comparison, direct correlation that is estimated between the genetic effects for a pair of cohorts has been proposed to estimate overlapping samples^26,27^, but it is confounded with genetic architecture, such as heritability underlying (**Table 1**). When there was heritability, the estimated correlation between genetic effects was biased and leads to incorrect overlapping samples for a pair of cohorts; when there was no heritability, the estimated correlation was correct and agreed well with the one estimated with *λ_meta_*. As existence of heritability is one of the reasons that trigger GWAMA, so *λ_meta_* is much proper in estimating overlapping samples between cohorts.

**Table 1.** The estimated correlation for a pair of cohorts via their summary statistics

| $n_1$ | $n_2$ | $n_{1,2}$ | $h^2$ | $M$ | $Q$ | $\gamma_{1,2} = \frac{n_{1,2}}{\sqrt{n_1 n_2}}$ | $\hat{\rho}_{1,2} \pm \text{SD}$ | $\hat{\gamma}_{1,2} \pm \text{SD}$ |
| --- | --- | --- | --- | --- | --- | --- | --- | --- |
| 1,000 | 1,000 | 100 | 0.25 | 30,000 | 1,000 | 0.1 | 0.1072±0.0064 | 0.101±0.0093 |
| 1,000 | 2,000 | 100 | 0.25 | 30,000 | 1,000 | 0.0707 | 0.0814±0.0054 | 0.0709±0.0088 |
| 1,000 | 5,000 | 100 | 0.25 | 30,000 | 1,000 | 0.0447 | 0.0615±0.0055 | 0.0425±0.0096 |
| 1,000 | 10,000 | 100 | 0.25 | 30,000 | 1,000 | 0.0316 | 0.0556±0.0063 | 0.0325±0.0099 |
| 1,000 | 1,000 | 1 | 0.25 | 30,000 | 1,000 | 0.001 | 0.0092±0.0056 | 0.0017±0.0093 |
| 1,000 | 2,000 | 1 | 0.25 | 30,000 | 1,000 | 0.0007 | 0.0126±0.0053 | 0.0006±0.0079 |
| 1,000 | 5,000 | 1 | 0.25 | 30,000 | 1,000 | 0.000447 | 0.0189±0.0060 | 0.0016±0.0090 |
| 1,000 | 10,000 | 1 | 0.25 | 30,000 | 1,000 | 0.000316 | 0.0259±0.0059 | 0.0008±0.0092 |
| 1,000 | 1,000 | 100 | 0 | 30,000 | 1,000 | 0.1 | 0.0996±0.0052 | 0.094±0.0085 |
| 1,000 | 2,000 | 100 | 0 | 30,000 | 1,000 | 0.0707 | 0.0704±0.0048 | 0.0712±0.0097 |
| 1,000 | 5,000 | 100 | 0 | 30,000 | 1,000 | 0.0447 | 0.0453±0.0057 | 0.0441±0.0090 |
| 1,000 | 10,000 | 100 | 0 | 30,000 | 1,000 | 0.0316 | 0.0335±0.0057 | 0.0325±0.0079 |

\* Q is the number of QTLs among M simulated loci. We also tried $Q = 100$ , the results were nearly identical.
$\gamma_{1,2}$ represents the true correlation due to overlapping samples
$\hat{\rho}_{1,2}$ represents the estimated correlation estimated via the method proposed by Bolormaa et al<sup>26</sup>, and Zhu et al<sup>27</sup>
$\hat{\gamma}_{1,2}$ represents the estimated correlation estimate via $\lambda_{meta}$ , $\hat{\gamma}_{1,2} = \frac{1 - \hat{\lambda}_{meta}}{\frac{2\sqrt{n_1 n_2}}{n_1 + n_2}}$ .

**Figure 8.**
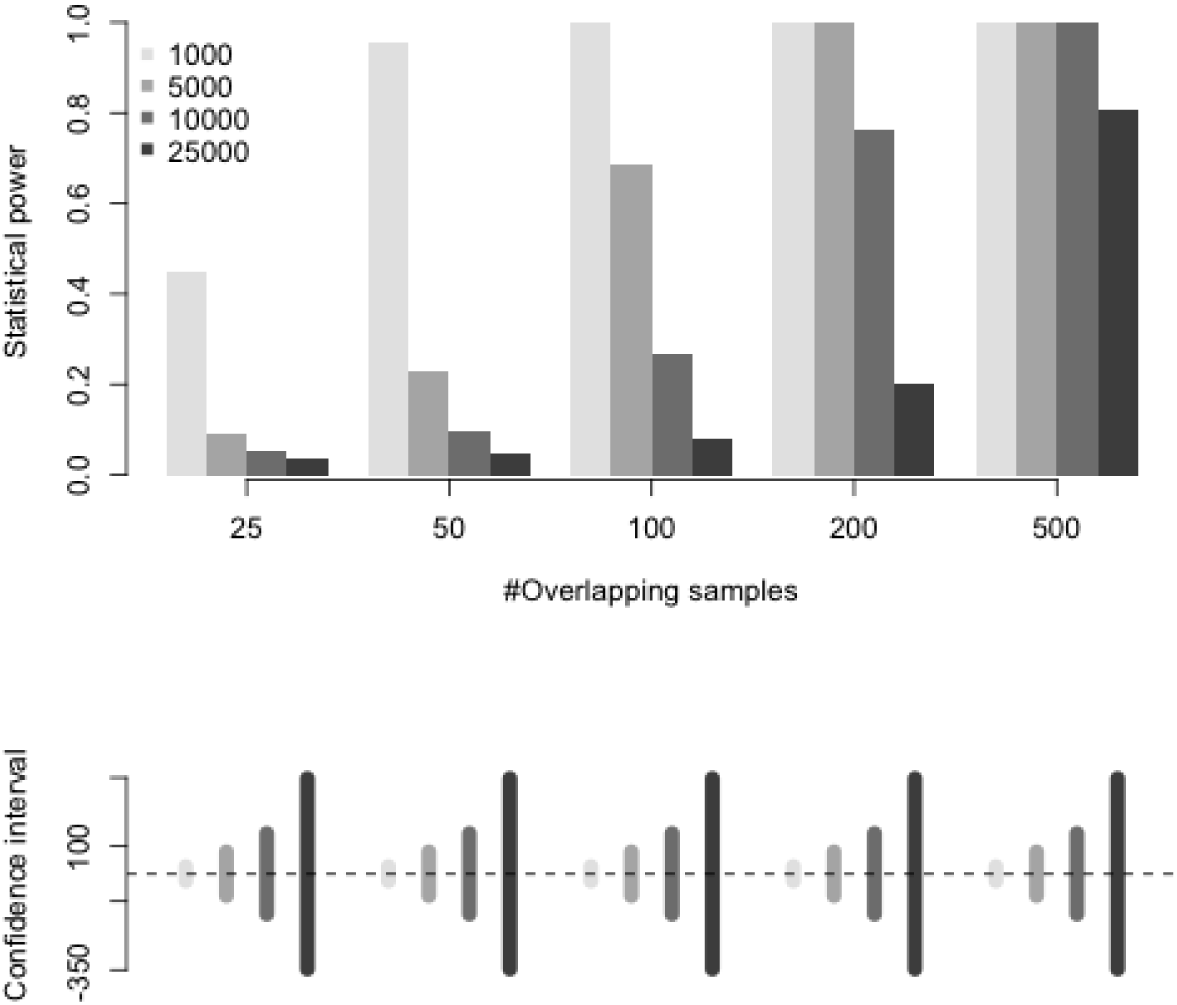
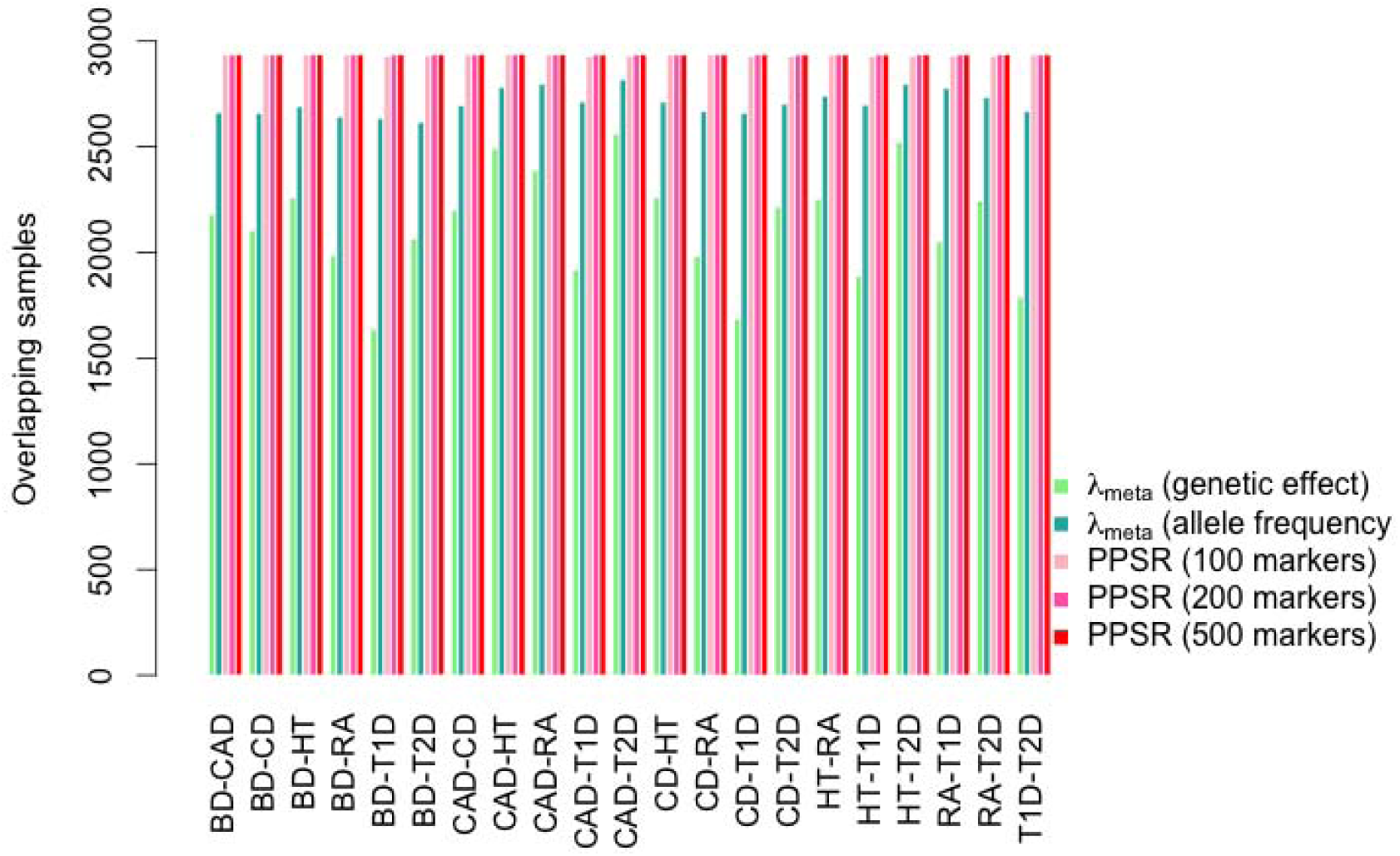
Pseudo profile score regression for the WTCCC 7 diseases. a) Statistical power for detecting overlapping samples between a pair of cohorts given type I error rate of 0.05. Top panel: The y-axis represents statistical power, and the x-axis the number of overlapping samples. Cohort 1 has 1,000, 5,000, 10,000, or 25,000 samples, and cohort 2 has 1,000 samples. The two cohorts have 25, 50, 100, 200, and 500 overlapping samples. Bottom panel: the corresponding 95% confidence interval is given for each scenario in the top panel. The statistical power is maximized when the two cohorts have the same sample size. b) Each cluster represents a pair of cohorts as denoted on the x-axis. Within each cluster, from left to right, the detected overlapping controls using *λ_meta_* based either on effect size estimates or minor allele frequency (MAF), PPRS using 100, 200, and 500 markers. WTCCC cohort codes: BD for bipolar disorder, CAD for coronary artery disease, CD for Crohn’s disease, HT for hypertension, RA for rheumatoid arthritis, T1D for type 1 diabetes, T2D for type 2 diabetes.

Another parameterization of *λ_meta_* is to estimate it from differences in allele frequencies between a pair of cohorts instead of differences between estimated effect sizes (**Supplementary notes**). We show that *λ_meta_* constructed on reported allele frequencies from genotyped loci from summary statistics can detect overlapping samples between two cohorts regardless of whether the GWAS is from quantitative traits or case-control data, even for pairs of different traits (**Supplementary notes and Supplementary notes**). For example, 2,934 common controls were shared across the WTCCC 7 diseases^7^. From the 21 pairwise *λ_meta_*, we estimated a mean of the number of overlapping samples, assuming overlapping controls only (**Supplementary notes**), of 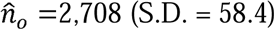, which was very close estimate to the actual number of overlapping samples (**Supplementary Fig. 8**). When constructing *λ_meta_* on the reported genetic effects and their sampling variance, the estimated mean estimate of the number of shared controls was 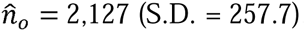, lower than that estimated from allele frequencies, which is likely due to real genetic heterogeneity between diseases (**Supplementary Fig. 8**). In practice, publically available summary statistics may not include sample specific allele frequencies, but may only be available with reference sample frequencies as a conservative strategy to prevent identification of individuals in a cohort.

### Detection of overlapping samples using pseudo profile score regression

GWAMAs have grown in sample size and in the number of cohorts that participate, and this trend is likely to continue. The probability that a sample is represented in more than one meta-analysis study is also likely to increase, in particular when very large cohorts such as UK Biobank and 23andMe provide data to multiple studies. While the metric *λ_meta_* can be transformed to give an estimate of *n_o_* between cohorts for quantitative traits, it cannot give an estimate of overlapping samples in case-control studies due to the ratio of the cases and controls in each study (**Supplementary notes**). Sharing individual genotype data (or imputed genotypes) across the entire study would make it easy to detect identical or near-identical genotype samples (representing real duplicate samples from individuals who participated in the two studies or monozygotic twins). In fact, only a small number of common SNPs is needed to detect sample overlap, and if this is known then individuals could be removed and summary statistics regenerated or the meta-analysis analysis itself can be adapted to correct for potential correlation due to *n_o_^28^*. However, in many circumstances, individual cohorts are not permitted to share individual-level data, either by national law or by local ethical review board conditions. To get around this problem, Turchin and Hirshhorn^10^ created a software tool, Gencrypt, which utilizes a security protocol known as one-way cryptographic hashes to allow overlapping participants to be identified without sharing individual-level data. To our knowledge, this encryption method has yet to be employed in meta-analysis studies. We propose an alternative approach, pseudo profile score regression (PPSR), which involves sharing of weighted linear combinations of SNP genotypes with the central meta-analysis hub. In essence, multiple random profile scores are generated for each individual in each cohort, using SNP weights supplied by the analysis hub, and the resulting scores are provided back to the analysis hub. PPSR works through three steps (**Supplementary notes and Supplementary Fig. 9**), and the purpose of PPSR is to estimate a relationship-like matrix of *n_i_* × *n_j_* dimension for a pair of cohorts, which have *n_i_* and *n_j_* individuals respectively. Each entry of the matrix is filled with genetic similarity for a pair of samples from each of the two cohorts, estimated via the PPSR.

**Figure 9.**
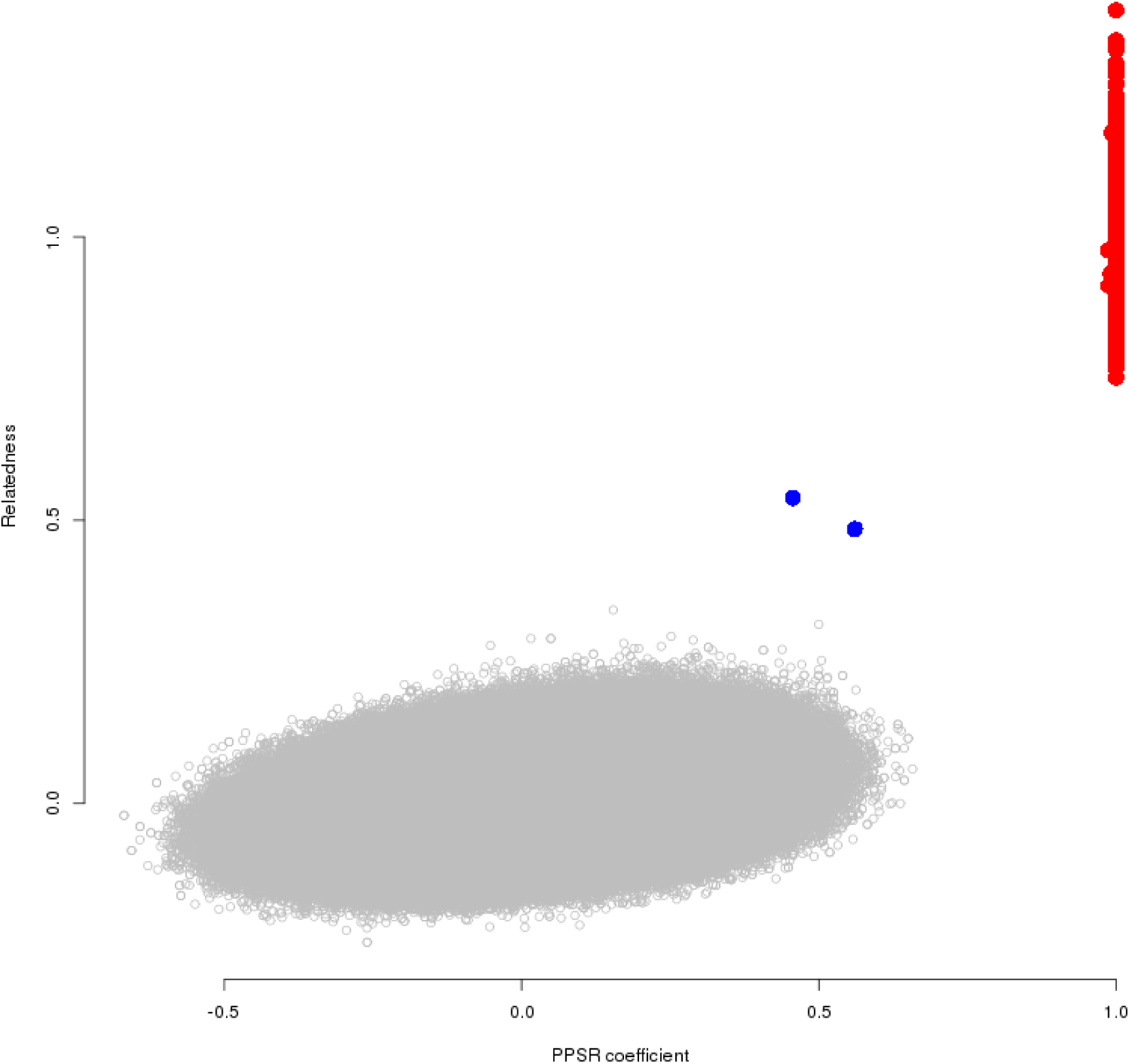
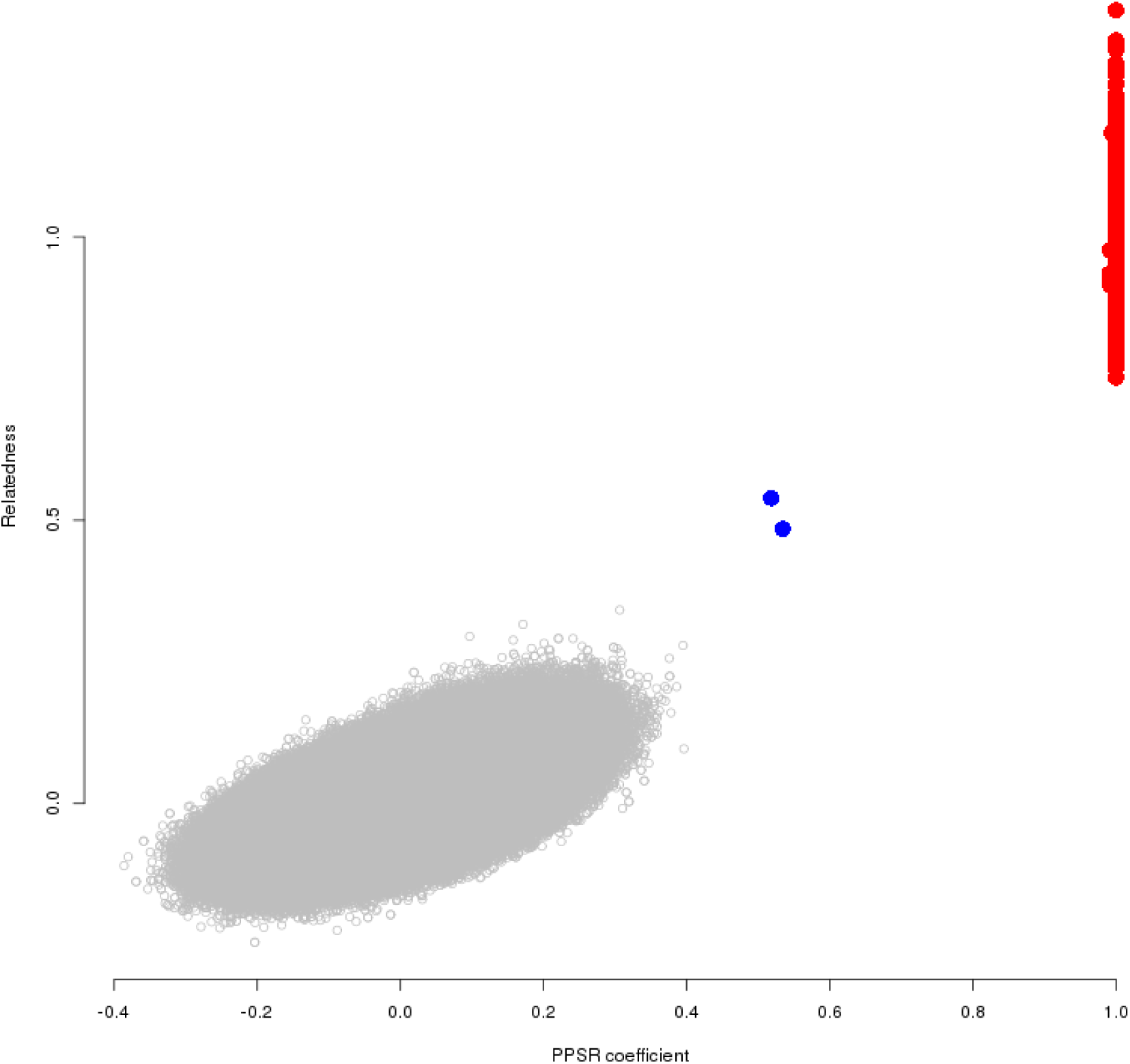
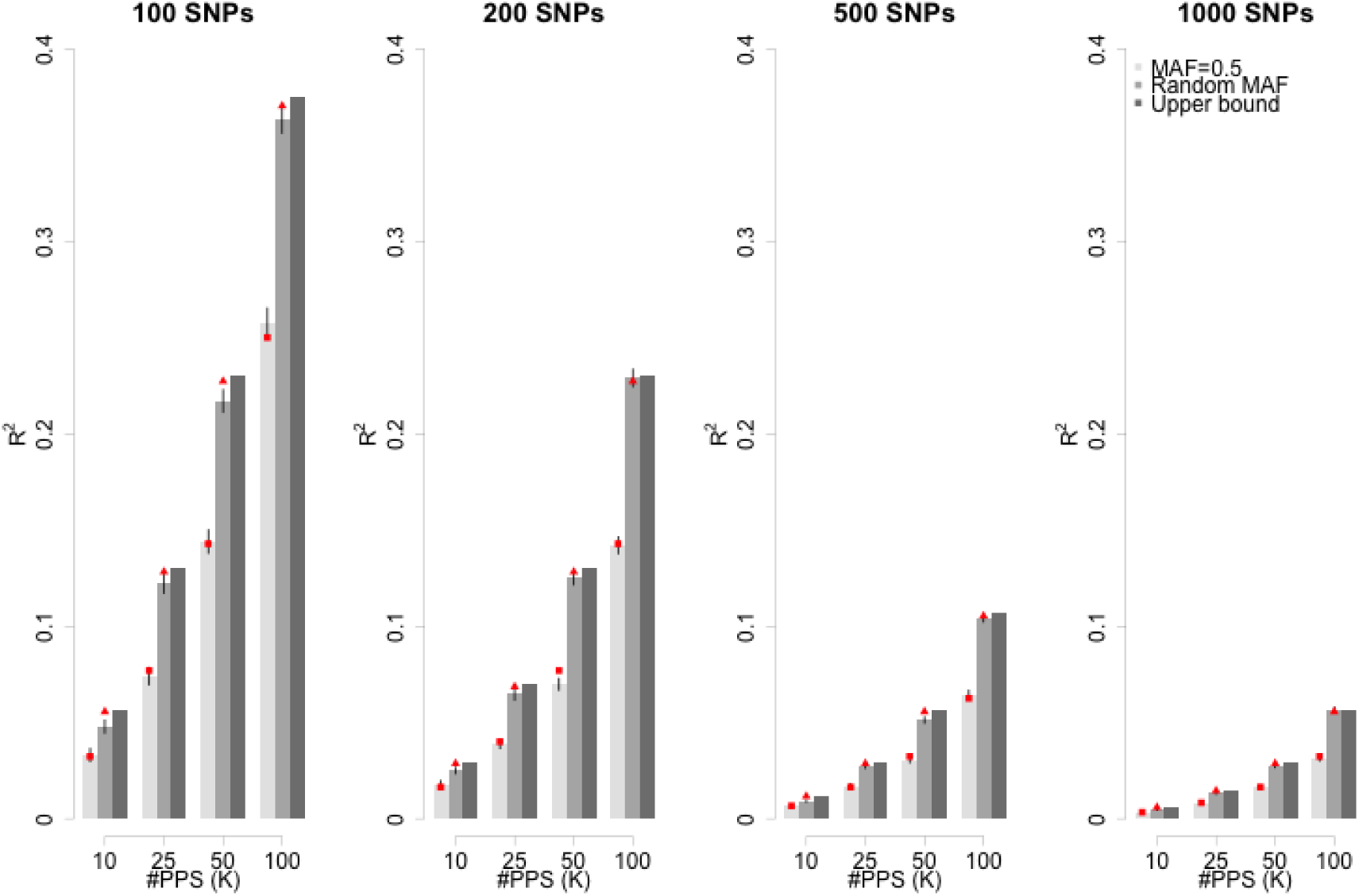
PPSR coefficients for identifying shared controls/relatives between WTCCC BD and CAD cohorts. **(a)** Illustration for regression coefficients between WTCCC BD and CAD from 57 pseudo profile scores (PPS) generated from 500 markers. The x-axis is the PPSR regression coefficients and y-axis is real genetic relatedness (as calculated from individual level genotype data). The red points are the shared controls between two cohorts, and blue points are first-degree relatives. **(b)** The PPS regression coefficients for detecting overlapping first-degree relatives using 286 PPS generated from 500 markers. **(c)** Decoding genotypes from the PPS. Given the set of profile scores, one may run a GWAS-like analysis to infer the genotypes. The ratio between the number of markers (*M*) and number of pseudo profile scores (*K*) determines the potential discovery of individual-level information. The higher the ratio and, the higher the allele frequency, the less information can be recovered. From left to right, the profile scores generated using different number of markers. The y-axis is a *R^2^* metric representing the accuracy between the inferred genotypes and the real genotypes. From left to right panels 100, 200, 500, and 1000 SNPs were used to generate 10, 20, 50, and 1000 profiles scores. In each cluster, the three bars are inferred accuracy using different MAF spectrum alleles, given with the SE of the mean.

We use WTCCC data as an illustration to detect 2,934 shared controls between any two of the diseases by PPSR. Among 330K unambiguous SNPs, which are not palindromic (A/T or G/C alleles), we randomly picked *M* = 100, 200, and 500 SNPs, to generate pseudo profile scores. It generated 21 cohort-pair comparisons, leading to the summation for 488,587,090 total individual-pair tests. To have an experiment-wise type I error rate = 0.01, type II error rate = 0.05 (power = 0.95) for detecting overlapping individuals, we needed to generated at least 57 pseudo profile scores (PPS). We generated scores *S =* [*s*_1_, *s*_2_*, s*_3_,…, *s*_57_], where each is a vector of *M* elements, sampled from a standard normal distribution (**Supplementary notes**). *S* is shared across 7 cohorts for generating pseudo-profile scores for each individual. In total 57 PPS were generated for each individual in each cohort. For a pair of cohorts, PPSR was conducted for each possible pair of individuals for any two cohorts over the generated pseudo-profile scores. Once the regression coefficient (*b*) was greater than the threshold, here *b =* 0.95, the pair of individuals was inferred to be having highly similar genotypes, implying that the individual was included in both cohorts (**Supplementary notes**).

When using 200 and 500 random SNPs, all the known 2,934 shared controls were detected from 21 cohort-pair-wise comparison; when using 100 randomly SNPs, on average 2,931 shared samples were identified, which is more accurate than using *λ_meta_* constructed using either genetic effects or allele frequencies (**Fig. 8b**). In addition, for detected overlapping samples, there were no false positives observed – consistent with simulations that show the method was conservative in the controlling type I error rate (**Supplementary notes**). For comparison, we also used the Gencrypt to detect overlapping samples using the same set of SNPs as used in PPSR. Although Gencrypt guidelines suggest use of at least 20,000 random SNPs^10^, selecting 500 random SNPs in the WTCCC cohorts also provided good accuracy with Gencrypt, and on average about 2,920 (99.6% of the shared controls) overlapping samples were detected, only slightly lower than PPSR. For example, for BP and CAD, Gencrypt detected 2,912 shared controls, but was unable to identify about 20 overlapping controls, due to missing data (on average 1% missing rate). Increasing the number of SNPs when using Gencrypt is likely to overcome the problem of missing data.

Furthermore, PPSR is able to detect pairs of relatives. For example, between the BD and CAD cohorts, two pairs of apparent first-degree relatives were detected (**Fig. 9a**). In order to find additional first-degree relatives between BD and CAD cohorts, at least 265 PPS were required to have a type I error rate of 0.01 and type II error rate of 0.05 (**Supplementary notes**) for a regression coefficient cutoff of 0.45, a threshold for first-degree relatives. As expected, all other individuals that did not show high relatedness did not reach the threshold of 0.45 of the PPS regression coefficient for first-degree relatives (**Fig. 9b**). Gencrypt did not detect any first-degree relatives.

The speed of PPSR depends on *n_i_* × *n_j_*, the sample sizes for a pair of cohorts, and the number of PPS for each cohort; for the WTCCC data there are 21 cohort-pair comparisons, and each pair took about 20 minutes, on a computer with a 2.3 GHz CPU, given about 5,000 × 5,000 = 25,000,000 comparisons. The average sample size of GIANT is about 1,500, and takes about 2 minutes for each pair of cohorts. The two largest datasets are deCODE with 26,790 samples and WGHS with 23,100 samples, and PPSR to detect overlapping samples takes about 8.5 hours. As each pair of individuals is computationally an independent unit, analysis jobs can be parallelized on a cluster. Therefore, even for meta-analyses involving many large cohorts, the computation time is not a limiting factor.

PPSR for each individual uses very little personal information and can be minimized so that there is very low probability of decoding it. One way to attempt to decode the genotypes from PPS is to reverse the PPSR, so that the individual genotypes can be predicted in the regression (**Supplementary notes**). The individual-level genotypic information that can be recovered by an analyst, who knows the matrix (the weights for generating PPS), is determined by the ratio between the number of markers (*M*) that generated PPS and the number of PPS (*K*). Therefore, inferred information on individual genotypes can be minimized and tailored to any specific ethics requirements. We suggest 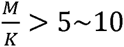 to protect the privacy with sufficient accuracy (**Fig 9c**). Of note, if a meta-analysis is conducted within a research consortium, the application of PPSR is even safer because the exchange of information is between the consortium analysis hub and each cohort independently.

## Discussion

In this study, we provide a set of metrics for monitoring and improving the quality of large-scale GWAMA based on summary statistics. These tools not only enrich the toolkit to analysts for GWAMA, but also provide informative summary and visualization for readers to understand the experimental design of GWAMA. As far as we know, no GWAMA to date has checked cohort-level outliers based upon population differentiation metrics or utilized estimated allelic effect sizes to identify and quantify sample overlap.

Using the *F_st_* derived genetic distance measure, we can place all cohorts on an inferred geographic map and can easily identify cohorts that are genetic outliers or that have unexpected ancestry. In application, we should note that the *F_st_* measure can identify unusual summary information, such as detected in the MIGEN cohorts from GIANT Consortium GWAMAs, in which the same allele frequencies were reported for all cohorts. Meta-PCA can also be used to infer the genetic background of cohorts. The high concordance between *F_pc_* and meta-PCA indicates the both methods are robust. In practice, mete-PCA may be much easier to implement when there are many cohorts, such as GIANT height cohorts and Metabochip BMI cohorts, but the coordinates of a cohort may be slightly shifted with inclusion or exclusion of other cohorts.

There are limitation for both *F_pc_* and meta-PCA. Firstly, the inference depends on the choice of reference cohorts. Meta-PCA is further upon the inclusion or exclusion of other cohorts. However, given the application of the data, we believe the impact will not influence the inference of the genetic background of cohorts in meta-analysis. Secondly, various mechanisms can give the identical projection in PCA^14^. The purpose of both methods is to find the discordance between demographic information and genetic information, or outliers. The projection is not attempt to discover the detailed demographic past that shapes a cohort.

Our third metric *λ_meta_* provides information on sample overlap and heterogeneity between cohorts by utilizing the estimated allelic effect sizes and their standard errors. In most meta-analyses, the overall *λ_meta_* is likely to be slightly greater than 1 solely due to unknown heterogeneity, slight as observed, in generating the phenotype and genotype data that cannot be accounted for by QC. The observed mean of *λ_meta_* for the GIANT height GWAMA was 1.03 but with more variation than expected by chance. The strong correlation between *λ_GC_* and *λ_meta_* indicated the reported sampling of the reported data were systematically driven by analysis protocols. For cohorts with *λ_GC_* < 1 and *λ_meta_* < 1, it is likely that the GWAS modeling strategy employed for GWAS in the cohort was too conservative, for example MIGEN cohorts might have on average too small sample size for each cohort. Conversely, for cohorts with *λ_GC_* > 1 and *λ_meta_* > 1 results are too heterogeneous, perhaps reflecting systematically smaller sampling variances of the reported genetic effects. As the GWAMA often uses inverse-variance-weighted meta-analysis^25^, such cohorts may lead to incorrect weights to the different cohorts in the meta-analysis, suggesting that the statistical analysis in meta-analyses can be improved by applying better weighting factors.

It is well-recognised that overlapping samples may inflate the type-I error rate of GWAMA and therefore lead to false positives. Although post-hoc correction of the test statistic is possible^26–28^, stringent quality control ruling out overlapping samples makes the whole analysis easier and lowers the risk of false positives. A better solution would be to rule out shared samples at the start, for pairs of cohorts that show deflated *λ_meta_*, and we propose PPSR to accomplish this.

In summary, to maximize the inference from multi-cohort GWAMA, accurate cohort-level information on allele frequencies, estimated effect sizes, and their sampling variance can be exploited to perform additional measures that are likely to lead to reduction in the number of false positives and increasing statistical power for gene discovery. All methods proposed are implemented in freely available software GEAR.

## Acknowledgements

This work was funded by Australian National Health and Medical Research Council Project and Fellowship grants (1011506, 613601, 613602, 1078901, 1078037), grants GM 099568 from the National Institutes of Health and the Sylvia & Charles Viertel Charitable Foundation. This study makes use of data generated by the Wellcome Trust Case-Control Consortium. A full list of the investigators who contributed to the generation of the data is available from www.wtccc.org.uk. We also thank from High Performing Computing support from the Information Technology group at the Queensland Brain Institute, The University of Queensland.

## Author contributions

GBC and PMV designed the study. GBC, PMV and SHL derived the analytical results. GBC performed all analysis. CGB and ZXZ developed the software. GBC and PMV wrote the first draft of the paper. MRR, JY, NW discussed results and methods, and provided comments that improved earlier versions of the manuscript. Other authors provided cohort-level summary statistics and contributed to improving the study and manuscript.

## Web resources

GEAR (GEnetic Analysis Repository): http://www.complextraitgenomics.com/

PGC: http://www.med.unc.edu/pgc/results

1000 Genomes Project: http://www.1000genomes.org/

---

Figure 1 Recovery of cohort-level genetic background and inference of their geographic locations for GIANT BMI Metabochip cohorts using the *F_st_* derived genetic distance measure.

Figure 2 Using the genetic distance spectrum to infer the geographic origins for GIANT height GWAS cohorts.

Figure 3 Comparison between Meta-PCA and genotype PCA on 1KG.

Figure 4 Recovery of cohort-level genetic background for GIANT BMI Metabochip cohorts using meta-PCA.

Figure 5 The recovery of cohort-level genetic background using meta-PCA analysis for GWAS height cohorts.

Figure 6 *λ_meta_* for the GIANT height GWAS cohorts.

Figure 7 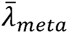 and *λ_gc_* for GIANT height GWAS cohorts.

Figure 8 Pseudo profile score regression for the WTCCC 7 diseases.

Figure 9 PPSR coefficients for identifying shared controls/relatives between WTCCC BD and CAD cohorts.

Table 1 The estimated correlation for a pair of cohorts via their summary statistics

## Method I: *F_st_* derived genetic distance

*F_st_* is a measure of genetic differentiation between populations. It is usually estimated using individual-level genotype data from multiple samples in two or more populations^1^. Here, we calculate *F_st_* using summary data on allele frequencies, which implicitly assumes Hardy-Weinberg equilibrium genotype frequencies within populations. We use summary statistic calculated *F_st_* as a metric for quality control for each cohort. If the allele frequencies reported for a cohort depart genome-wide from its expectation based on known ancestry due to technical artifacts, then we may observe an unexpected *F_st_* value when comparing to a reference panel of know ancestry.

We calculate *F_st_* between each cohort and a reference panel, choosing the appropriate reference sample depending on the purpose of the analysis. For the inference of global-level diversity, we chose YRI, CHB, and CEU as the reference panels. For the inference of within-Europe diversity, we chose CEU, FIN, and TSI as the reference panels. As the different allele frequencies across three samples reflected the real diversity among these reference panels, we did not apply any exclusion criteria on the reference allele frequency. Nevertheless, as GIANT height GWAS samples were imputed to the HapMap panel, the majority of SNPs matched to the 1KG reference samples comprised common SNPs. After ranking the calculated *F_st_* in ascending order for all matched SNPs, we sampled 30,000 *F_st_* evenly along the ordered *F_st_*. These 30,000 markers are quasi-independent and evenly distributed across the genomes. The mean of the 30,000 *F_st_* was employed to represent the *F_st_* measure between a cohort and a reference panel. The sampled 30,000 markers may differ from one pair of cohorts to another pair, but as tested resample 30,000 markers caused ignorable changes of the mean of *F_st_*. Another reason we chose 30,000 markers is that there are around 30,000 quasi-independent markers for GWAS data as observed in empirical data and expected from theory^2,3^.

In this study, *F_st_* is calculated from the allele frequencies estimated from cohorts, provided as summary statistics. *F_st_* is treated as a data statistic for measuring allele frequency differentiation. In general the interpretation of *F_st_* can vary with context^4^.

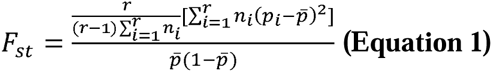
with *p_i_* the estimated reference allele frequency in population *i* from a sample of *n_i_* alleles, 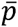 is the weighted average frequency in the entire sample, and *r* is the number of populations. Here, we only compared each cohort to the 1KG reference panel, so *r = 2* and the equation becomes

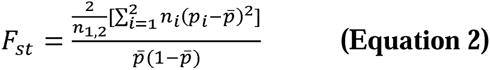
in which *n*_1,2_ = *n*_1_ + *n*_2_, and 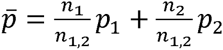 is the mean allele frequency. Alternative estimators for *F_st_* are possible, and a comprehensive comparison of different *F_st_* estimators was recently reported^5^.

If the two cohorts are not that different in terms of their allele frequencies, for example, the cohorts from European nations, 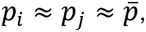,

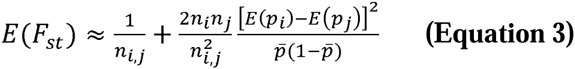

At the right side of the equation, the first term represents the sampling variance for allele frequency for a pair of cohorts, and the second term represents the allele frequency difference due to divergence from a common ancestor. The estimated *F_st_* is influenced by sample size, and 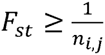, which is the sampling variance of *F_st_* for a pair of cohorts^1^. As each 1KG reference population has a sample size around 100, there is no disproportionate impact of sample size in calculating *F_st_*.

*F_st_* **Cartographer algorithm**. The purpose of using the *F_st_* Cartographer algorithm is to find the coordinates of a cohort given its *F_st_* to the reference populations. The algorithm can be expressed in Cartesian geometry. Given three reference populations, a target cohort has three *F_st_* measures, *F*_1_, *F*_2_, and *F*_3_, respectively. Given a Cartesian coordinate system, the coordinate for these three reference populations are (*a*_1_, *b*_1_), (*a*_2_, *b*_2_), and (*a*_3_, *b*_3_), respectively. The algorithm tries to find the coordinates 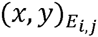 on each the edge (*E*_i,_*_j_*) hat connects reference populations *i* and *j*

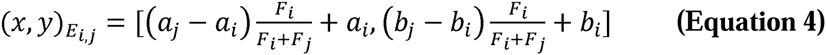

The coordinates of the gravity of triangle, (*x*, *y*)*_G_*, that connects *E*_1,2_, *E*_1,3_, and *E*_2,3_ are

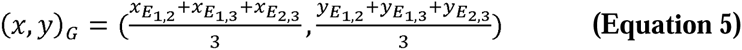

**Inference of cohort origins at the global level**. To assess genetic background, for each cohort we calculated its *F_st_* values using CEU, CHB, and YRI as the reference panel, respectively. We denote these three *F_st_* values as *F_CEU_*, *F_CHB_*, and *F_YRI_*. These values reflect genetic distances between a cohort and the reference panels - the greater the value the further the genetic distance. We developed an algorithm called *F_st_* cartographer, which can map a cohort to global genetic variation as previously observed using individual level data from principal component analysis^6^. The steps in the algorithm are as follows (**Supplementary Fig. 1**):

Create the coordinates for the reference samples. Without loss of generality, these three reference populations form an equilateral triangle, and we set the length of each edge to unity. For example, the coordinates CEU, CHB, and YRI are 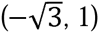, 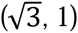, and (0, −2), respectively, and connecting the coordinates of the three reference populations formed an equilateral triangle - the reference space. The gravity of this equilateral triangle is the origin of the Cartesian space. The choice for the coordinates for the reference population is arbitrary.

**Step 1 Create a cohort triangle using Equation 4**. Finding a point the distances of that to both ends, which represent two populations, is proportional to the ratio of the *F_st_* values of the cohort to these two reference populations. Similarly, find the points on the other two edges. For example, Finland Twin Cohort (FTC) had *F_CEU_ =* 0.0102, *F_YRI_* = 0.153, and *F_CHB_* = 0.099. On the CEU-YRI edge, a point split the length to 0.0102:0.153, was 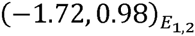; On the CEU-CHB edge into 0.0102:0.099, was 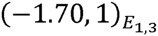; and on the YRI-CHB edge into 0.153:0.099, was 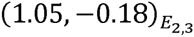. Connecting the three coordinates created a “FTC” triangle inside the reference triangle.

**Step 2 Find the gravity of the cohort triangle using Equation 5**. The gravity of the “FTC” triangle had its coordinates of 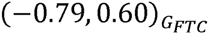, which is inferred as the geographic coordinates for FTC in *F_PC_* space. It had relative distances of 1.03, 2.55, and 2.72 to CEU, CHB, and YRI, respectively. The shorter the distance, the closer the genetic background is.

**Step 3 Repeat Steps 1, and 2 until the gravity of each cohort is found**.

Plots of the coordinates for each cohort will show the relative distance of each cohort to the reference samples. If a cohort has equal distances to three reference populations, its gravity will be close to the origin of the reference triangle.

## Method II: Principal component analysis for cohort-level allele frequencies

PCA has been widely used in genetics^7^ and recently proposed for controlling population stratification for GWAS^8,9^. We provide a new method that uses cohort-level allele frequencies, often provided as summary statistics in meta-analysis. We call the new method as meta-PCA.

Meta-PCA is based on a *G =* (*C + K*) × *M* matrix, which includes *K* reference populations and *C* cohorts of question on *M* markers. In *G*, the *m^th^* column represents the reported reference allele frequencies for the *m^th^* marker for (K+C) cohorts. The kernel correlation matrix for PCA is constructed on 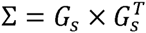, in which *G_s_* is the standardization for for *G* for each locus (on each column of *G*). Compared with individual-level data PCA, in the context of meta-PCA each cohort can be viewed as an individual in the conventional sense. Given Σ matrix, all the implementation is the same as the individual-data PCA.

There are efforts in establishing genetic interpretation for PCA ^8,10–12^ The interpretation of meta-PCA could be approached by *F_pc_* as described in the last section.

## Method III: The detection of overlapping samples with *λ_meta_*

**Inference of cohort origins at the within-Europe level**. To assess genetic background, for each cohort we calculated its *F_st_* values using CEU, FIN, and TSI as the reference panel, with coordinates 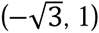, 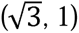, and (0, −2), respectively. For FTC, it had *F_st_* values of 0.0102, 0.0052, and 0.0157, to CEU, FIN, and TSI, respectively. Using the *F_st_* Cartographer algorithm, the gravity of the FTC triangle had its coordinates of 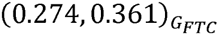. It had relative distances of 2.10, 1.59, and 2.42, to CEU, FIN, and TSI, respectively.

**Genealogical subspace**. Furthermore, we partition the *F_PC_* space into three subspaces. For example, given coordinates of 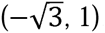, 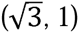, and (0, −2), for CEU, FIN, and TSI, respectively, connecting the origin and the coordinates for any two reference populations created a subspace, which is defined as a genealogical subspace. We had three genealogical subspaces: CEU-FIN genealogical subspace, CEU-TSI genealogical subspace, and FIN-TSI genealogical subspace, respectively. If a cohort is located inside a subspace, it indicates that this cohort may be derived from these two reference populations that creates the genealogical subspace.

For European cohorts, the coordinates calculated from *F_st_* Cartographer algorithm mirror the origins of geographic locations of the cohorts, similar, but less refined, to what has been observed in previous studies using individual level data for European samples^13,14^.

**Effective number of overlapping samples (*n*_o_)**. If a pair of cohorts has overlapping samples, it leads to a correlation of the estimated genetic effects for each locus. In the recent literature, two kinds of correlation due to overlapping samples were introduced. The first one was defined by directly calculating correlation between all estimated test statistics, *r = cor*(*Z*_1_, Z_2_), in which *Z* is a vector of *M* matched loci between two cohorts^15,16^. The second one was defined on the correlation for single locus given overlapping samples, as introduced by Lin and Sullivan^17^. We used the second definition, and then extended the correlation due to any relatives, a generalization of Lin and Sullivan.

For a pair of cohorts of sample sizes *n*_1_ and *n*_2_ (*n*_1_ ≥ *n*_2_), for *M* matched loci which have GWAS summary statistics, for example additive effects and their standard errors. For the *m^th^* locus, estimated association effect sizes are *b*_1_*_.m_* and *b*_2_*._m_* with sampling variance 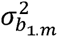 and 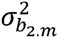, respectively. *b*_1_ is assumed to be drawn from a normal distribution 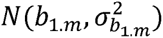, and 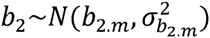. In cohort 1, 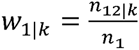 is proportion of samples with a *k^th^*-degree relatives in cohort 2 with *n*_12\_*_k_* the number of relatives of kth degree relatives shared between the samples; the phenotypic variance is assumed to be the same across the cohorts for a quantitative trait. For a locus, the genetic effect is estimated by linear regression *y*_1_ *= a* + *b*_1_*x* + *e* in cohort 1 (the index for the locus is dropped for convenience). If the sampling variance of a locus is assumed to be the same for any subset of samples 

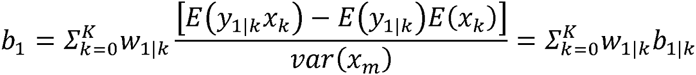

The standard error of *b_m_* is 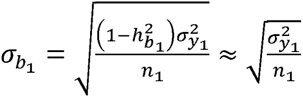, in which 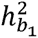 is the proportion of phenotypic variance explained by the locus and 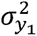 is the phenotypic variance of the trait. The sampling variance for 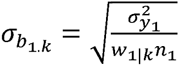. This decomposition of the genetic effect can be applied to cohort 2. Consequently, the covariance between *b*_1_ and *b*_2_ for the locus is

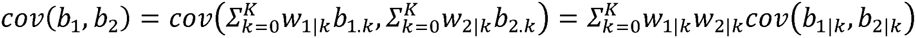
in which 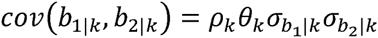 is the covariance between the genetic effects estimated in two cohorts due to the *k*-degree relatives. *ρ_k_* is the phenotypic correlation for the *k*-degree relatives, and *θ_k_* is the genetic relatedness for the *k*-degree relatives. 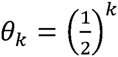 is the coefficient of identity for descent. For duplicated samples, *ρ*_0_ *= h^2^* + *ρ_e_*_|0_, in which *h^2^* is the heritability, and *ρ_e_*_|0_, the environmental correlation to be close 1 for overlapping samples; for other relatives (*k* ≥ 1), *ρ_e_*_|0_ ≈ *θ_k_h*^2^.

**Correlation between the estimated genetic effects**. The covariance can be generalized as 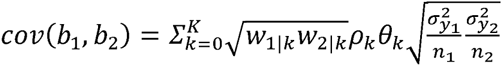. After adjustment by the sampling variance, the correlation between *b*_1_ and *b*_2_ is 

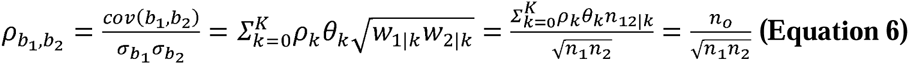
in which 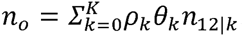, is the effective number of overlapping samples averaged over all relative pairs that are across the two cohorts. As the variance explained by each locus is small, and after further weighted by *θ_k_*, the contribution from overlapping relative is small. When ignoring the first and higher degree relatives *n_o_* equals the contribution from overlapping samples. This is consistent with the results from Lin and Sullivan^17^, who considered overlapping samples only. So, the correlation at any single locus is largely determined by the overlapping samples (*n*_12|0_) for summary statistics.

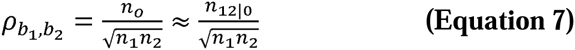

So, in the text hereafter, *n_e_* indicates overlapping samples only, otherwise specified.

**Correlation for case-control studies**. The theory above is based on a quantitative trait, but it holds approximately true for case-control studies if a locus is from the null distribution of no association with the disease. Given *n*_12_.*_ctrl_* overlapping controls and *n*_12_.*_cs_* overlapping cases, for a locus associated with disease its correlation of the regression coefficient is 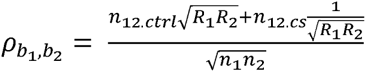 as indicated by Lin and Sullivan^17^, in which *R_i_* is the ratio between cases and controls in the *i^th^* cohort. When it is balanced case-control design – *R. =* 1, 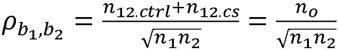 resembles the correlation for quantitative traits. However, it should be noticed that for case control data, *n_e_* is confounded with the number of overlapping cases and controls.

**Theory for** *λ_meta_*. For the summary statistics between a pair of cohorts for the *m^th^* locus, we can construct a statistic

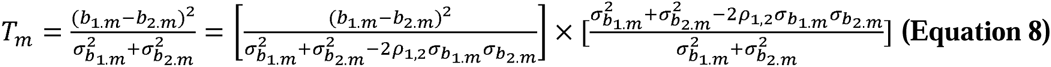
in which *ρ*_1,2_ is the correlation between *b*_1,_*_m_* and *b*_2,_*_m_*.

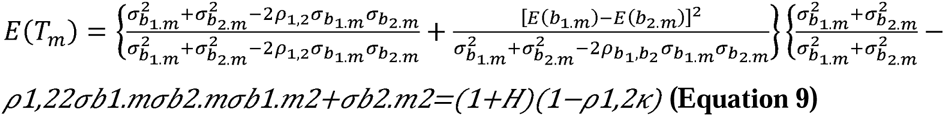
in which 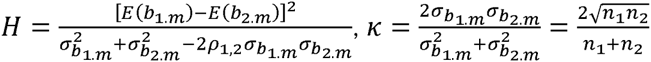, and 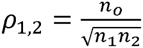, as defined in Equation 12, is the correlation for this locus due to overlapping samples between this pair of cohorts. Of note, *ρ*_1,2_ is same for each locus regardless of a null locus or a locus associated to genetic effects. For convenience, the subscript *b* was dropped in the text hereafter.

Under the null hypothesis of no heterogeneity (*H =* 0) and no correlation (*ρ*_1,2_ = 0), 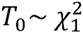, a standard 1-degree-of-freedom chi-square distribution. 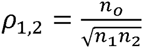, in which *n_o_* is the effective number of overlapping samples. Of note, since the majority of markers are likely sampled from the null distribution or have very small effect sizes, we can approximate *E*(*b*_1_*_.m_*) *=* 0 and *E*(*b*_2_*_.m_*) = 0, and therefore *H ≈* 0 for most marker pairs between a pair of cohorts. For the *m^th^* marker that is in linkage disequilibrium with causal variants, 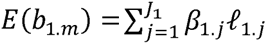, in which *J*_1_ is the number of causal variants in linkage disequilibrium with the *m^th^* marker for cohort 1, *β*_1_*_.j_* is the *j^th^* causal variants in linkage disequilibrium with the *m^th^* marker, and *ℓ*_1_*_.j_* is the LD correlation between the *m^th^* marker and the *j^th^* causal variant^18^. Similarly for 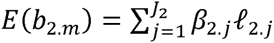. If the cohorts are from the same ethnicity, the difference in the LD correlation can be ignored, for example for samples from cohorts with European ancestry. So, under a polygenic model *H* is expected to be zero, or close to zero.

The *T* statistic is calculated for each matched SNP between a pair of cohorts. After ordering all *T* values, we evenly sample 30,000 independent markers from the order statistic of all *T* values. Each pair of cohorts may sample *T* values based on 30,000 markers different from another pair of cohorts.

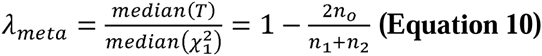
in which 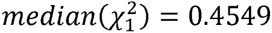. Under the null hypothesis of no heterogeneity and overlapping samples (*n_o_* = 0), plotting the ordered *T* against its corresponding quartiles from 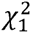, will be along the diagonal, leading to *λ_meta_* = 1. Heterogeneity between two cohorts, equivalent to a “negative” number of overlapping samples, will drive *λ_meta_* > 1, and overlapping samples will make *λ_meta_* < 1. The distribution of *λ_meta_* can be assessed via the beta distribution, and *λ_meta_* follows asymptotically a normal distribution *N*(1,0.0136) given 30,000 independent markers.

**Factors that influence** *λ_meta_*. A number of factors will influence the *λ_meta_*. 1) Sample overlap, including close relatives across cohorts, reduces the value of *λ_meta_* (**Supplementary Fig. 2**) Conservative modeling, such as inclusion of covariates in the association model that are genetically correlated with the phenotype or the ‘genomic control’ approach (adjusting the sampling variance with 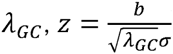), will inflate the sampling variance, and deflate *λ_meta_*. 3) Genetic heterogeneity, which can be caused by differences in genetic architecture or methodological difference, will inflate *λ_meta_*. 4) As characterized by Equation 10, the lower bound (cohort 2 is completed included in cohort 1, given *n*_1_ > *n*_2_ = *n*_0_) of *λ_meta_* is 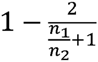, upon the ratio of the samples sizes of the two cohorts.

**Estimating overlapping samples**. As shown in Equation 10, *λ_meta_* is a linear function of *n_o_*, hence the statistical power to detect overlapping samples is equivalent to asking how *λ_meta_* departs from the null distribution. Assuming *H =* 0, the overlapping samples can be estimated as 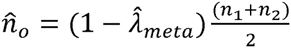, and 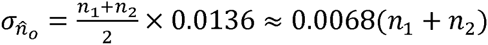 given 30,000 independent markers. Hence, using summary statistics only the proportion of overlapping samples can be estimated for quantitative traits. Given the type I error rate of 0.05 (*α* = 0.05), the statistical power for detecting 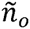 overlapping samples between two cohorts is 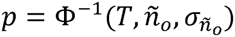, in which Φ^−1^ represents the accumulation power function of a normal distribution with the mean of *ñ_o_* and standard deviation of 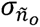. The statistical power is determined by *T*, the threshold for significance, *ñ*_0_, the real overlapping samples, and 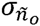, the standard deviation of the null hypothesis that there is no overlapping samples. Without loss of generality, 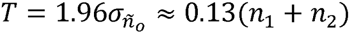 given *α =* 0.05. The 95% confidence interval is [−0.13(*n*_1_ + *n*_2_), 0.13(*n*_1_ + *n*_2_)]. The statistical power is maximized when *n*_1_ = *n*_2_, i.e. when a pair of cohorts has the same sample size.

For case-control studies, as 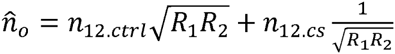, the estimate cannot distinguish between overlapping cases and overlapping controls; when *R*_1_ = 1 and *R*_2_ = 1 (balanced case-control design for both cohorts), 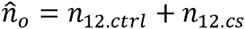, indicating the overall overlapping samples between two cohorts, summed across cases and controls. If we know that only controls (cases) were shared between two cohorts, then 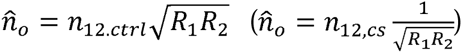, so then an estimate of *n_o_* indicates the number of overlapping controls (cases). Therefore, quantifying overlapping samples for case-control studies is more difficult than that for quantitative traits.

## Method IV: Pseudo profile score regression (PPSR)

**PPSR** resembles the previously proposed Gencrypt method^19^, but PPSR is more powerful in detecting various degree of relatives and more robust to missing data and imputation errors. For each individual, the PPS can be generated as below

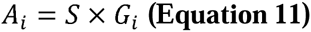
in which *A_i_* is the PPS for the *i^th^* individual, *S* is a *K × M* score matrix, and *G_i_* is vector for the genotypes for the chosen *M* loci.

In detail, 

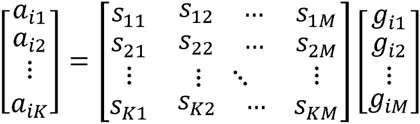
in which *a_ik_* is the *k^th^* profile score for the *i^th^* individual, *s_km_* is the additive effect at the *m^th^* locus (*m* from 1 to *M*) for the *k^th^* profile score, and *g_im_* is the standardized genotype at the *k^th^* locus for the *i^th^* individual. Each *s*, the pseudo genetic effect, follows a standard normal distribution *N*(0,1); each pseudo genetic effect is independent to another. For each PPS, 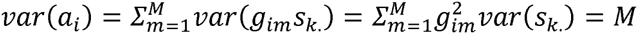, in which *s_k_* is the *k^th^* column for the *S* matrix, and on average each locus explains 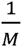 of the variation. For an individual a pair of PPS, say *a_l_*_1_ and *a_l_*_2_, has 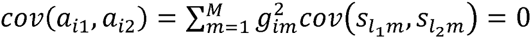.

Each PPS can be seen as a trait with *h^2^ =* 1 because it does not have any sampling variance. For a pair of individuals, individual *i* and individual *j*, when both *A_i_* and *A_j_* have been standardized, their covariance for the *k^th^* PPS (*a_i_*_1_, *a_j_*_1_) = *θh*^2^ in which *θ* is the relatedness scores in terms of identity by state^20^. Depending on the relatedness between a pair of individuals, *θ* = 1 for monozygous twins or to a duplicated sample, *θ* = 0.5 for first-degree relatives such as parent and offspring or full sibs. In general, for *r^th^*-degree of relatives, *E*(*θ_r_*) = 0.5*^r^*.

The theory presented above provides a theoretical basis for detecting overlapping samples using PPS other than sharing individual level genotypes. Assuming that each individual has *K* independent PPS (*A_i_* having *K* elements), for individual *i* and *j*, we can regress *A_i_* on *A_j_*,

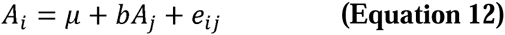
in which *μ* is the grand mean, *b* is the regression coefficient, and *e_ij_* is the residual. 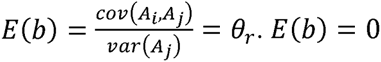 if individual *i* is not correlated with individual *j, E*(*b*) = 0.5 for first-degree relatives, and *E*(*b*) *=* 1 if individual *i* and *j* are genetically same, say an overlapping sample or the homozygous twins. The sampling variance of *b* is 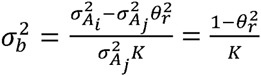. Under the null distribution for no related or overlapping samples, 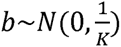. The residual *e_ij_* accounts the discordant genotypes, including missing genotypes and genotyping or imputation errors. For current GWAS data, after quality control, the discordant rate is often smaller than 1%.

If now we have *C* cohorts for which the individual genotypes of which cannot be disclosed to the central analysis hub, overlap between cohorts can be identified if PPS are supplied. By regressing their PPS to each other the overlapping individuals could be detected if *b* ≈ *θ_r_*. Assuming there are *N_c_* samples in each cohort, a total of 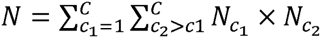 regressions need to be carried out as defined in Equation 12. If we want to control the experiment-wise type I error rate *α* under the null hypothesis and type II error rate *β* (with power= 1 − *β*) for *b = θ_r_*, the required number of pseudo profile scores for each individual is

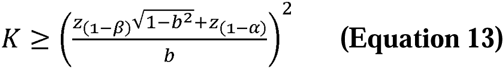
in which and *z*_(1 −_*_β_*_)_ are *z*_(1 −_ *_α_*_)_ are *z* scores under the given *p*-values at the subscripts. To accommodate technical errors, such as missing genotypes and genotype error, a cutoff of 0.95 for *b* is adopted for detection of overlapping samples, and 0.4~0.45 for detecting first-degree relative.

The standardization of genotypes can either use the allele frequency from each cohort, or from a reference sample. Throughout the study, we used the allele frequency calculated from WTCCC bipolar disorder cohort as the reference, and using it as an approximation to standardize genotypes for all cohorts in comparison.

**Workflow for PPSR**. Given the statistical method for detecting overlapping samples as described above, the whole workflow for detecting can be split into three steps (**Supplementary Fig. 9**).

**In step 1, the required type I and type II error rates are defined and from that the required number of pseudo profiles to be generated**. The GWAMA central analyst selects consensus SNP markers across cohorts, and determines additive effects matrix *S* that will be used to generate pseudo profile scores for each cohort. In order to avoid strand issues, the loci having palindromic loci (A/T alleles or G/C alleles) are excluded.

**In step 2, each cohort generates PPSR for each individual with the set of consensus markers and the marker weights received from the GWAMA coordinator**. After generate the PPS, they send them back to the coordinator. This will be a file that contains N rows and K columns with pseudo-profile scores.

**In step 3, the coordinator runs PPSR for each sample in a cohort on each PPS generated for another cohort**. The final product of running PPSR is to generate a *n_i_* × *n_j_* matrix for a pair of cohorts, which have *n_i_* and *n_j_* samples respectively. For each pair of individuals in comparison, we take the one from cohort *i* as the response variable and from cohort *j* as the predictor variable in PPSR. In principle, swapping the response variable and the predictor variable do not affect the performance of PPSR. Each entry, the regression coefficient of PPSR, in the *n_i_* × *n_j_* matrix represents genetic similarity for these pair of individuals in comparison. Once the regression coefficients are above the threshold, it indicates there are samples duplicated. The central analyst can then request each cohort that is implicated in containing samples that are also in other cohorts to drop those samples, without revealing where the duplication occurred.

**Privacy issues when using PPSR**. As the exchange of the PPS is within a meta-analysis facility, it is not as vulnerable as that of releasing the GWAS summary to the public domain as discussed in previous studies^21–23^. However, as PPS are generated from genotypes, it is worth to consider whether the PPS will reveal individual genotype information, or can be decoded from PPS. As a demonstration for the principle-of-proof, we consider to reverse Equation 11 to estimate genotypes. We consider the case where the additive effect matrix in Equation 11 is known, otherwise it is nearly impossible to recover genotype information. Given the workflow of PPSR, the analysts who coordinate the meta-analysis know the additive effect matrix, *S* in Equation 11, and receive PPS from each cohort have the information to decode genotypes that are employed to generate PPS.

After reversing Equation 11, using the standard regression method, the genotype in each locus can be estimated as 

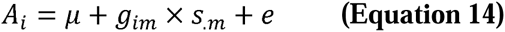
In detail,

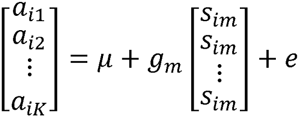
in which *s_m_* is the *m^th^* column in the additive effects matrix in Equation 11. Although *E*(*g_im_*) = *g_im_*, which is an unbiased estimate of the genotype, its sampling variance is 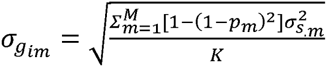. The sampling variance can be further written as 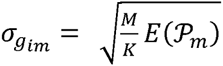 because 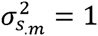 and [1 − (1 − *p_m_*)^2^] is denoted as *p_m_*. The greater the ratio between 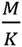 and *E*(*p_m_*), the larger the sampling variance, and consequently the lower probability to construct the real genotype.

Without loss of generality, the accuracy of the estimated 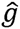, a continuous variable, and *g*, a discrete variable with values of 2, 1, and 0, can be measure using the squared correlation (*R*^2^)^24^, 

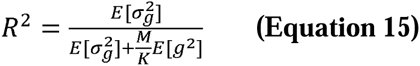
in which *E*(*g*^2^) and 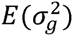 are:

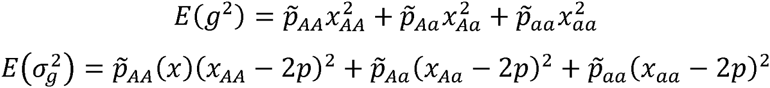
*x_AA_* = 2, *x_Aa_* = 1, and *x_aa_* = 0 if *A* is the reference allele, and 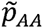, 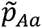, and 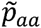 are weighted frequency given the distribution of *g*. 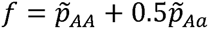.

When the reference allele frequency follows a uniform distribution between (*a*_1_, *a*_2_), assuming that the loci follow Hardy-Weinberg proportions, *p_AA_* = *p*^2^, *p_Aa_* = 2*pq*, and *p_aa_*= *q*^2^, in which *p* follows a uniform distribution between *a*_1_ and *a*_2_ and *q =* 1 − *p*.

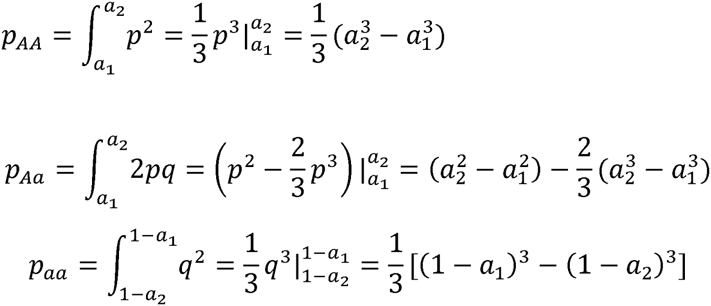
and 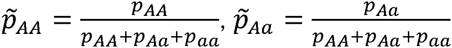, and 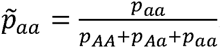.

If the reference allele frequency follows a uniform distribution between (0, 0.5), 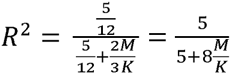.

Given *M* loci with MAF of 0.5, the expected frequencies for *AA, Aa*, and *aa* are 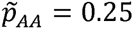, 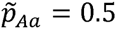, 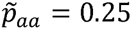, and *f =* 0.5. *E*(*g^2^*) *=* 1.5, and 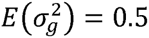. Plugging them in to the Equation 13 leads to 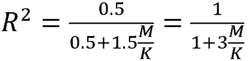.

Equation 13 can be rewritten as 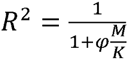, in which *φ* = 3 if MAF is 0.5, and *φ* = 1.6 if MAF in nearly from a uniform distribution. From Equation 13, it is easy to calculate the ratio between the number of markers and the number of PPS given a controlled *R*^2^,

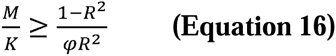

For uniform distribution of MAF, if *R^2^ ≤* 0.1 is set as the threshold, 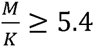; if *R^2^ ≤* 0.05,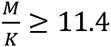 and if 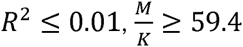. In general, the higher the ratio between *M* and *K*, the less information can be inferred. We suggest 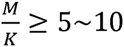 may be sufficient.

